# Multivariate pattern analysis of fMRI data for imaginary and real colours in grapheme-colour synaesthesia

**DOI:** 10.1101/214809

**Authors:** Mathieu J. Ruiz, Michel Dojat, Jean-Michel Hupé

## Abstract

Grapheme-colour synaesthesia is a subjective phenomenon related to perception and imagination, in which some people involuntarily but systematically associate specific, idiosyncratic colours to achromatic letters or digits. Its investigation is relevant to unravel the neural correlates of colour perception in isolation from low-level neural processing of spectral components, as well as the neural correlates of imagination by being able to reliably trigger imaginary colour experiences. However, functional MRI studies using univariate analyses failed to provide univocal evidence of the activation of the ‘colour network’ by synaesthesia. Applying Multivariate (multivoxel) Pattern Analysis (MVPA) on 20 synaesthetes and 20 control participants, we tested whether the neural processing of real colours (concentric rings) and synaesthetic colours (black graphemes) shared patterns of activations. Region of interest analyses in retinotopically and anatomically defined visual regions revealed neither evidence of shared circuits for real and synaesthetic colour processing, nor processing difference between synaesthetes and controls. We also found no correlation with individual experiences, characterised by measuring the strength of synaesthetic associations. The whole brain, searchlight, analysis led to similar results. We conclude that identifying the neural correlates of the synaesthetic experience of colours may still be beyond the reach of present technology and data analysis techniques.

## Introduction

Synaesthesia is a subjective experience shared by only a fraction of the population (Simner et al., 2006; Chun & Hupé, 2013; Simner & Carmichael, 2015; Rouw & Scholte, 2016; Watson et al., 2017), offering, in principle, an opportunity to study the neural bases of subjective experience, drawing on individual differences just like in neuropsychology, but involving healthy people. Moreover, colour, the typical prototype of a qualia (what it feels like to perceive something) is the most often cited (or at least studied: Ward, 2013) content of the synaesthetic experience. However, the very subjective nature of the synaesthetic experience represents a major obstacle when trying to set an objective and operational definition, as required in an experimental protocol. Not only subjective descriptions may vary a lot between subjects (Flournoy, 1893), but also within subjects when asked to complete the same questionnaire again (Edquist, Rich, Brinkman, & Mattingley, 2006) or when describing their subjective experience of colour for different letters (Hupé, Bordier, & Dojat, 2012b). Using psychophysical tests, the synaesthetic experience of colour appears more similar to imagined or remembered than perceived colours (Witthoft & Winawer, 2013; Chiou & Rich, 2014; Hupé & Dojat, 2015; Janik McErlean & Banissy, 2017). The experience of synaesthetic colours can be indeed formally described as a form of mental imagery, since it occurs without any corresponding spectral stimulation. The obligatory experience of colour when exposed to letters or digits may therefore justify the label of ‘intrusive visual imagery’ (Reeder, 2016). Unfortunately, this simplification does not help much with defining the phenomenological content of synaesthesia, since self-reports of mental imagery show at least as much diversity as those of synaesthesia (Galton, 1880), with mixed evidence about whether the presence of synaesthesia may relate to individual differences in mental imagery (Chun & Hupé, 2016). One may, however, study how much synaesthesia requires the neural resources involved in visual perception. This bottom-up approach, which does not address the phenomenological issue, can at least be operationalized. Moreover, grapheme-colour synaesthesia offers a unique opportunity regarding the neural correlates of imagination as it restrains both individual variability and the content specificity of visual imagery. Last but not least, synaesthetic colours are systematically triggered by letters and digits, unlike “regular” mental imagery that depends on both the good will and the (uneven) ability of subjects.

Several brain imaging studies have compared activations in the visual cortex for real and synaesthetic colours, whose majority did not reveal any overlap. There were even questions whether activations triggered by synaesthetic stimuli, when observed, were in fact related to the synaesthetic experience at all (Hupé & Dojat, 2015). This surprising ‘Null’ result may be due to methodological limitations since only massive univariate analysis of brain imaging data were used so far, which may reveal only processes well localized in the brain (Hupé et al., 2012b). Multivariate (multivoxel) Pattern Analysis (MVPA) does not suffer from such a restriction. MVPA provides a way to reveal how information is encoded by the brain (Cox & Savoy, 2003; Norman, Polyn, Detre, & Haxby, 2006; Formisano & Kriegeskorte, 2012; Hebart & Baker, 2017). It has been applied successfully to the decoding of aspects of mental images (Thirion et al., 2006; Reddy, Tsuchiya, & Serre, 2010). Using fMRI, here we simply asked whether classifiers trained on patterns of blood oxygenation dependent signals (BOLD responses) elicited by different coloured stimuli could predict which synaesthetic colours were experienced by synaesthetes when seeing achromatic letters and digits. We studied in particular the early stages of visual processing by identifying cortical areas V1 to V4 in each subject, using retinotopic mapping, thus avoiding the problems related to structural normalization (Poldrack, 2007; Hupé, 2015). We also explored the whole visual cortex (including parts of the parietal cortex) using regions of interest based on a probabilistic atlas (Eickhoff et al., 2007), and performed whole brain searchlight analyses (Kriegeskorte, Goebel, & Bandettini, 2006). We compared all the measures obtained in synaesthetes with those obtained in a group of non-synaesthetes to take into account any potential non-specific effect related to the choice of stimuli. We also took into account the individual variability of the synaesthetic experience: without any possibility to characterize objectively the different phenomenological accounts, we measured the strength of the synaesthetic associations (Ruiz & Hupé, 2015).

## Materials and Methods

### Participants

We tested 20 synaesthetes and 20 non-synaesthetes. Synaesthetes (17 women) were between 21 and 42 years old (M = 27.9, SD = 5.5). Recruitment was diverse and opportunistic, based on self-referral following publicity on internet: lab webpage, *Facebook* event, announcements on university networks in Grenoble and Paris. Potential participants, after a first phone interview, were asked by email to fill-up a questionnaire to describe their synaesthetic associations and for grapheme colour associations to send us a list of those. Synaesthetes were included if they had a sufficient and diverse number of letter-colour and digit-colour associations as required by the design of our experiments (see below). When they came to the laboratory to perform the experiments, they had a semi-directed interview to evaluate the phenomenology of their synesthetic associations. They also ran a modified version of the “Synaesthesia Battery Test” (Eagleman, Kagan, Nelson, Sagaram, & Sarma, 2007) to choose precisely the colour of each letter and digit. This procedure was also used as a retest to confirm the validity of the first-person reports (Ruiz & Hupé, 2015): in all subjects, all chosen colours matched those indicated by print or by name in the questionnaire. In addition, objective measurements of synaesthetic associations were obtained by Stroop-like tests (see below in the ‘Protocol’ section). Seven of the included synaesthetes had already participated in psychophysics experiments between 2007 and 2010 (Ruiz & Hupé, 2015).

Control participants were recruited after synaesthetes to match their demographic statistics (16 women, age range between 23 and 38 years old, M = 28.5, SD = 4.3), following similar advertisement strategies as well as soliciting colleagues at the Grenoble Institute of Neuroscience. Interviews were conducted to verify the absence of any type of synaesthesia, not only the absence of grapheme-colour associations. We chose not to run any consistency score with the control subjects in order not to prime them to do any voluntary association between graphemes and colours before the tests in the scanner. It could be argued that some of the controls may have had implicit synaesthetic associations they were not aware of, as it sometimes happens. In any case, this unlikely possibility could not bias our results because most of our analyses did not require any direct comparison of the performances by synaesthetes and controls.

The study was performed in accordance with the Declaration of Helsinki, it received approval by the Institutional Review Board of Grenoble (CPP 12-CHUG-17, approval date 04/04/2012) and written, informed consent was obtained from all subjects. A medical doctor verified that all subjects were without past or current brain disease and had no detected cognitive deficit. All subjects had normal colour perception on the Lanthony D-15 desaturated colour test (Richmond products), and normal or corrected to normal eyesight (then using MRI-compatible glasses).

### Materials

Stimuli: for each synaesthete, we tried to identify four pairs of graphemes made of one letter and one digit that had similar colour associations. We never chose graphemes for which a synaesthete indicated several colours. We tried to find pairs of red, green, blue and yellow (R, G, B, Y) graphemes, but we were only partially successful and in some cases we selected a pair from the most saturated colours available. Figure 1 shows the actual letters and digits with colours used in the experiments. Only 13 subjects named the pairs red, green, blue and yellow; other colours were named orange, violet, fuchsia and brown, as well as light and dark blue or green. Syn08 and syn48 had a pair made of two letters. Since each synaesthete was tested with a different set of stimuli, each control subject was tested with the stimuli of a specific synaesthete (with the exception of syn10 who had no matched control, by mistake; two controls were tested instead with the stimuli of syn11. Paired comparisons were therefore based on 38 subjects).

**Figure 1.**
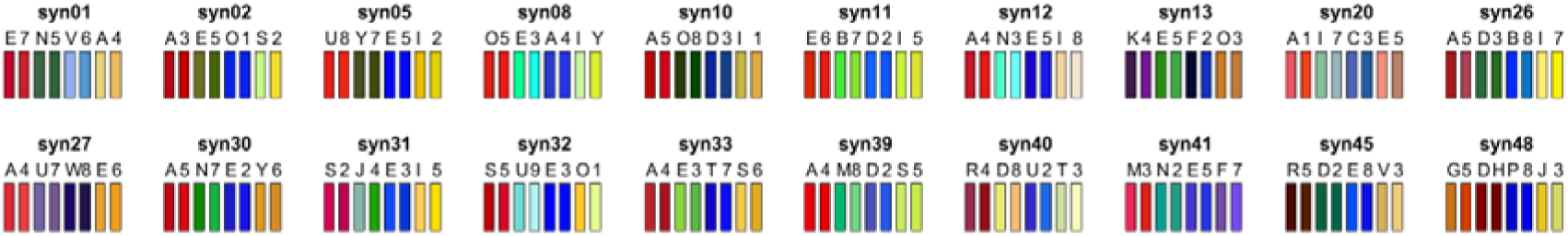
Letters and digits used for each synaesthete, with their corresponding synaesthetic RGB colours (the rendering of the colours using the projector in the scanner was different).

In the MR scanner, we presented these letters and digits in black at the centre of the screen (upper case, Helvetica font, extent up to 2 degrees eccentricity) over a grey uniform background (CIE xyY [0.29 0.3 77.4], half of the maximal luminance of the screen). Stimuli were projected on a translucent screen at the back of the scanner by a video projector Epson EMP 8200. We used a spectrophotometer (PhotoResearch PR 650) for colour and luminance measurements used to compute calibrated images. We also presented dynamic concentric rings (square luminance profile, similar to the stimuli used by Brouwer & Heeger, 2009, except for the absence of anti-aliasing so as to use only the colours selected by each synaesthete), with the exact same (real) colours as those chosen by each synaesthete for each grapheme. The choice of colours matching the individual grapheme ‘R, G, B, Y’ colour associations was done again by each synaesthete in the scanner over the same grey background, using a house-modified MRI compatible, comfortable, 10-button console controller, and the colour-picker of the “Synaesthesia Battery Test” as was done previously outside of the scanner. The same coloured rings were used for each matched control. The rings extent was also up to 2 degree eccentricity and the spatial frequency was 3 cycles/degree (six circles). The phase of the rings changed randomly at 6 Hz to almost nullify visual effects induced by the absence of anti-aliasing.

These stimuli were chosen with the purpose of training and testing classifiers (see below, “Data analysis: classifications”). Briefly, we wanted to use the BOLD responses to the coloured rings to train classifiers on colours, and the BOLD responses to graphemes to train classifiers on synaesthetic colours. This required choosing pairs of dissimilar graphemes, i.e. a letter and a digit, to try to avoid that the classifier trained on some shared form features, but rather on their common associated colour. This also implied that decoding should not be feasible that way based on the responses of control subjects. The use of pairs of graphemes also allowed the training on letters and testing on digits (or the reverse), with success in principle possible only for synaesthetes, based on their synaesthetic colour associations. The careful matching procedure of synaesthetic colours allowed the training of classifiers on real colours and testing on graphemes to identify which brain regions, if any, coded both real and synaesthetic colours. Again, any decoding success would in principle be possible only in synaesthetes.

Classifiers would be trained and tested on four categories, ‘R, G, B, and Y’, referring either to the real or the synaesthetic prototypical colours that we tried to select. To maximize classification performances, categories should be perceptually disjoint. Figure 2 represents the actual colours used in the scanner for each synaesthete within the CIE L*a*b* colour space, which is more perceptually uniform than the CIE xyY space. As was already obvious in Figure 1, differences of luminance were important to distinguish stimuli. Figure 2 illustrates that the colour and luminance distances were not similar across subjects between categories and within pairs, leading to unequal clusterisation. We could even expect some confusions by the classifiers for some subjects (e.g. “green/yellow” for syn11, “red/blue” for syn13 or “blue/yellow” for syn41). While the maximal theoretical performance achievable by classifiers was therefore below 100%, classifiers could however obtain more than the 25% chance performance in every subject. For all the analyses (described below), we tested if the performance of classifiers across subjects was correlated with the ratio of colour distance (indicated for each subject in Table 1) as measured in the L*a*b* space; we did not find any evidence of that, except for the classification of colours in the searchlight analysis, when testing the group of synaesthetes: we found one significant cluster (106 voxels, 2862 mm^3^), in the left fusiform gyrus, peaking at MNI XYZ = [−27 −73 −4], extending from about V4 to FG4, in line with the involvement of these regions in colour processing.

**Figure 2.**
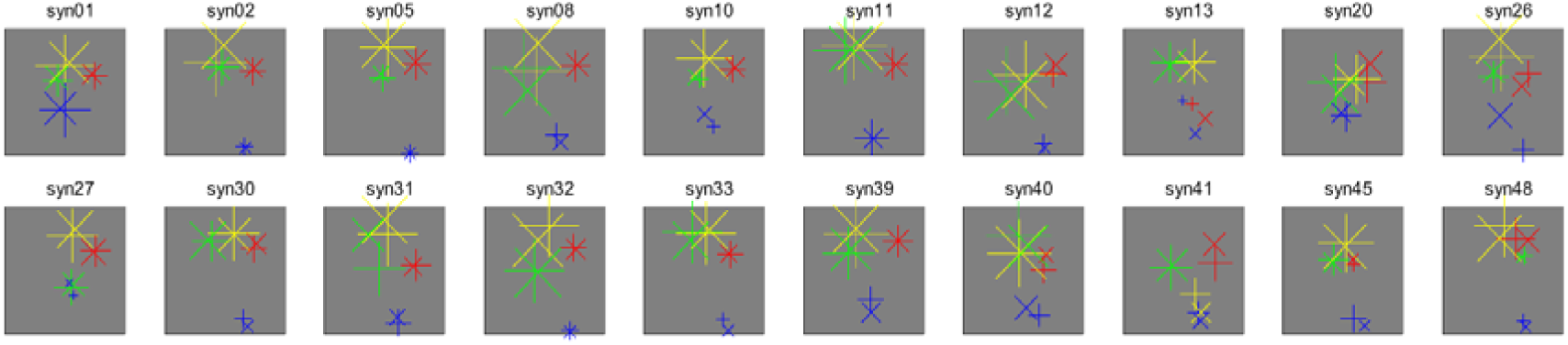
Colour coordinates in the CIE L*a*b* space of the stimuli used for each synaesthete, corresponding to the idiosyncratic synaesthetic colours of letters (+) and digits (x). The colours of the crosses are arbitrary and correspond to the four categories the classifiers had to distinguish. The size of the crosses is proportional to luminance (marker size = 0.4*L, where max(L) = 100; axes limits are +/- 130, possible range being −128 to +127).

**Table 1.**
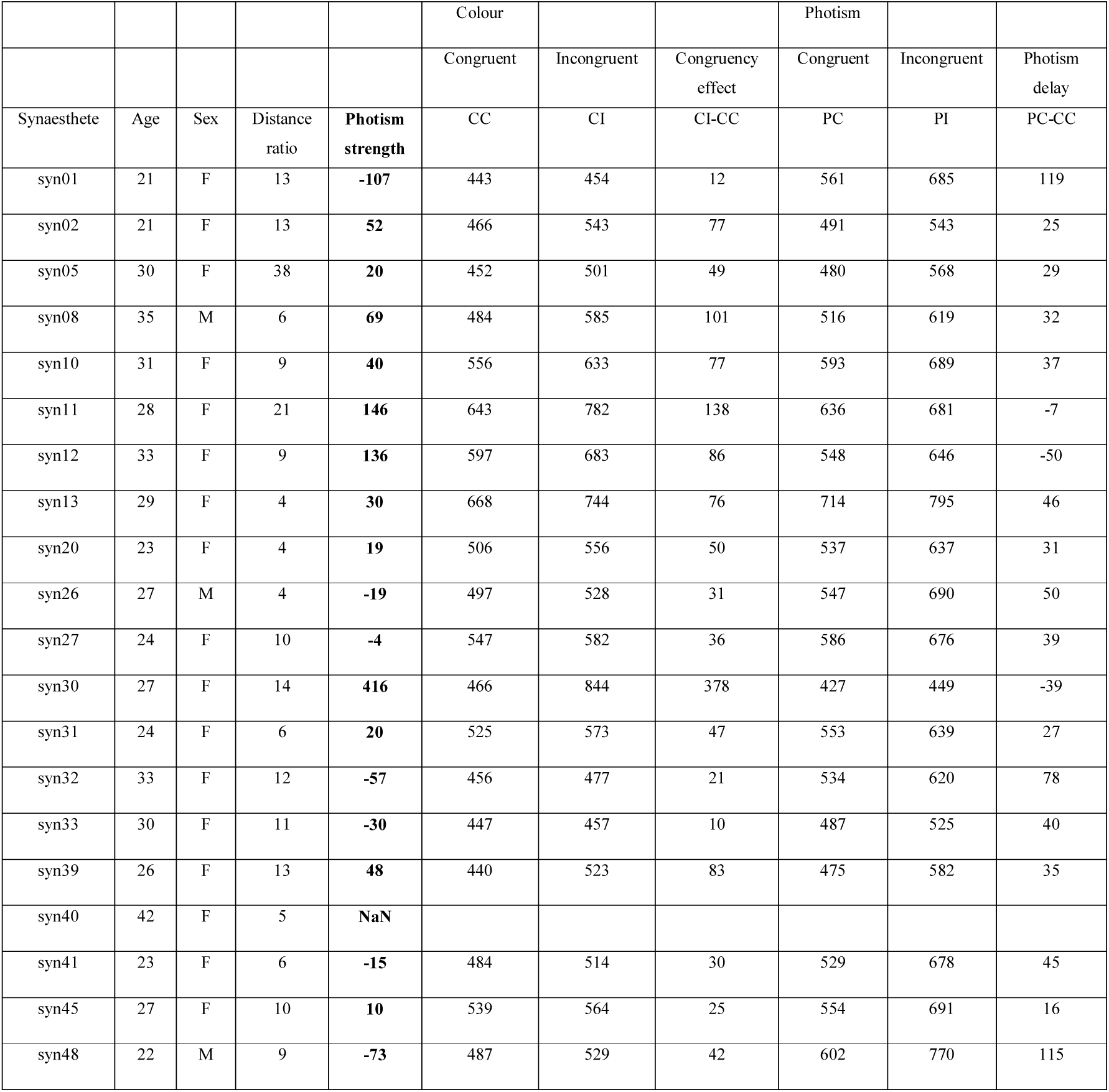
Demographics and characteristics of the tested synaesthetes. The variable ‘Distance ratio’ is a measure of clusterisation of the pairs of colour (average between-cluster distance divided by average within-cluster distance, measured in the L*a*b* space). The higher the value, the better the clusterisation (see Figure 2). The variable ‘Photism strength’ is the measure of the strength of synaesthetic associations as developed by Ruiz et al. (2015), based on the results of Stroop like tests. The next columns indicate the median time measured in *ms* for each subject to name either the real (‘Colour’) or the synaesthetic colour (‘Photism’) of the letters and numbers shown in Figure 1, when the real colour was either congruent or incongruent with the synaesthetic colour indicated by each synaesthete. The ‘Congruency effect’ was measured as the difference of response time in the ‘Colour’ condition. ‘Photism delay’ is the difference between naming the real and the synaesthetic colours, and ‘Photism strength’ is the difference of these two values. The interpretation of this index is only relative (the zero value does not have any special meaning). The data of syn40 data were not consistent (see text).

|  |  |  |  |  | Colour |  |  | Photism |  |  |
| --- | --- | --- | --- | --- | --- | --- | --- | --- | --- | --- |
|  |  |  |  |  | Congruent | Incongruent | Congruency effect | Congruent | Incongruent | Photism delay |
| Synaesthete | Age | Sex | Distance ratio | Photism strength | CC | CI | CI-CC | PC | PI | PC-CC |
| syn01 | 21 | F | 13 | **-107** | 443 | 454 | 12 | 561 | 685 | 119 |
| syn02 | 21 | F | 13 | **52** | 466 | 543 | 77 | 491 | 543 | 25 |
| syn05 | 30 | F | 38 | **20** | 452 | 501 | 49 | 480 | 568 | 29 |
| syn08 | 35 | M | 6 | **69** | 484 | 585 | 101 | 516 | 619 | 32 |
| syn10 | 31 | F | 9 | **40** | 556 | 633 | 77 | 593 | 689 | 37 |
| syn11 | 28 | F | 21 | **146** | 643 | 782 | 138 | 636 | 681 | -7 |
| syn12 | 33 | F | 9 | **136** | 597 | 683 | 86 | 548 | 646 | -50 |
| syn13 | 29 | F | 4 | **30** | 668 | 744 | 76 | 714 | 795 | 46 |
| syn20 | 23 | F | 4 | **19** | 506 | 556 | 50 | 537 | 637 | 31 |
| syn26 | 27 | M | 4 | **-19** | 497 | 528 | 31 | 547 | 690 | 50 |
| syn27 | 24 | F | 10 | **-4** | 547 | 582 | 36 | 586 | 676 | 39 |
| syn30 | 27 | F | 14 | **416** | 466 | 844 | 378 | 427 | 449 | -39 |
| syn31 | 24 | F | 6 | **20** | 525 | 573 | 47 | 553 | 639 | 27 |
| syn32 | 33 | F | 12 | **-57** | 456 | 477 | 21 | 534 | 620 | 78 |
| syn33 | 30 | F | 11 | **-30** | 447 | 457 | 10 | 487 | 525 | 40 |
| syn39 | 26 | F | 13 | **48** | 440 | 523 | 83 | 475 | 582 | 35 |
| syn40 | 42 | F | 5 | **NaN** |  |  |  |  |  |  |
| syn41 | 23 | F | 6 | **-15** | 484 | 514 | 30 | 529 | 678 | 45 |
| syn45 | 27 | F | 10 | **10** | 539 | 564 | 25 | 554 | 691 | 16 |
| syn48 | 22 | M | 9 | **-73** | 487 | 529 | 42 | 602 | 770 | 115 |

Note that luminance variations constitute a major difference, due to the constraint of using synaesthetic colours, with other MVPA studies of the neural correlates of colour processing, which used isoluminant stimuli (Brouwer & Heeger, 2009; Parkes, Marsman, Oxley, Goulermas, & Wuerger, 2009). We do not know how differences along the luminance axis should be perceptually scaled to differences along the green/red opponent colours a* axis and the blue/yellow opponent colours b* axis. This question is probably an ill-posed problem when studying brain correlates of colour perception, since at the cortical level visual circuits rely on both (but with different degrees) the parvo- and the magno-cellular pathways (Tootell & Nasr, 2017).

### Protocol

Each subject ran three fMRI sessions of about 1 hour. In addition, synaesthetes ran a 1 hour psychophysics experiment (before or interleaved with fMRI experiments, depending on schedule availability) to measure the strength of their synaesthetic associations using variants of Stroop tasks.

All the details of the psychophysics experiment as well as the results of 11 synaesthetes are published (Ruiz & Hupé, 2015). Briefly, eight graphemes (repeated 36 times each) were presented randomly either with the colour chosen by each synaesthete (congruent condition) or with the synaesthetic colour of the other presented graphemes (incongruent condition). Synaesthete had to name as fast as possible the real colour of the grapheme (‘colour’ task). Response times were measured *a posteriori* based on the audio recording. The procedure was then repeated, but synaesthetes had this time to name as fast of possible the name of the synaesthetic colour (called a photism) they associated to each grapheme, which was also either congruent or incongruent with the real colour of the stimulus (‘photism’ task). The index of the strength of synaesthetic associations (‘photism strength’) combined two measures: the response time difference for congruent and incongruent stimuli in the colour task, which reflects the difficulty to inhibit synaesthetic associations; the response time difference to name the real and the synaesthetic colours (in the congruent condition), which reflects how easily synaesthetes retrieve the synaesthetic colour.

The data of one synaesthete (syn40) could not be analysed because the chosen orange and yellow/green colours revealed too similar (see Figure 2) and were not named consistently over the course of the experiment. Table I provides a summary of the data. It shows that even synaesthetes who obtained a relatively low score of photism strength (e.g., syn01) were very fast at naming the synaesthetic colour of letters and numbers, even though the real colour of the stimulus was systematically varied. Moreover, they very rarely made any mistake (Ruiz & Hupé, 2015). Such a task would be extremely difficult to perform by any non-synaesthete trying to memorize (without training) random colour associations. These data therefore provide a further objective validation of the genuineness of the synaesthetic experience of these participants as well as an estimate of the strength of the associations.

### fMRI experiments

The MR experiments were performed at the IRMaGe MRI facility (Grenoble, France) with a 3T Philips Intera Achieva, using a 32 channels coil. The experiments can be decomposed successively in three “sessions” (about 1 hour each), “runs” (a few minutes), “blocks” (1 minute) and “events” (1 second). One session was dedicated to retinotopic mapping and functional localizer runs using pictures of objects, words and coloured stimuli (Mondrian). These latter runs were included to test whether voxels involved in the decoding of synaesthetic colours were located in regions well-defined functionally, respectively the Lateral Occipital Complex (LOC: Grill-Spector, Kourtzi, & Kanwisher, 2001), the Visual Word Form Area (VWFA: Dehaene & Cohen, 2011) and “colour centres” (Hupé et al., 2012b). We did not have the opportunity to use those localizers (see Results). Retinotopic mapping was performed strictly as described in a previous study (Bordier, Hupé, & Dojat, 2015), using the Brain Voyager analysis pipeline to define in each subject the ventral and dorsal as well as the left and right parts of areas V1, V2, V3 and V4 (ventral only). The parameters of the EPI functional images were TR/TE: 2000/30 ms, excitation pulse angle: 80°, acquisition matrix: 80×80, bandwidth: 54.3 Hz/pixel, isotropic nominal resolution: 3 mm, 30*2.75 mm thick slices with 0.25 mm interspace covering the whole visual cortex, with four additional dummy scans. To allow the precise alignment of functional scans across sessions, a high-resolution structural image of the brain was also acquired using a T1-weighted MP-RAGE sequence. The sequence parameters were TR/TE: 25/2.3 ms, excitation pulse angle: 9°, 180 sagittal slices of 256*240 (read x phase), bandwidth: 542.5 Hz/pixel isotropic nominal resolution: 1 mm, for a total measurement time of 4 min 31 s.

Another session was dedicated to the “synaesthesia” protocol (a structural image was also acquired with the same parameters as in the first session, in the middle of the functional runs). Twelve functional runs were acquired. The parameters of the EPI functional images were identical to those used for the retinotopic mapping experiment but TR: 2500 ms for an acquisition volume of 45 slices covering the entire brain with a total measurement time of 3 min 47 s. In each functional run, stimuli of one type only were presented: letters, digits, concentric rings with the synaesthetic colours of letters, or concentric rings with the synaesthetic colours of digits. The session contained three successive sequences of four runs, each run with a different stimulus type (with a different random order of stimulus type in each sequence). Each run contained 3*60 s blocks of a rapid-event paradigm, separated by 10 s fixations. Stimuli of different “colours” were presented pseudo-randomly in each block to optimize the estimation of the main effects. For example, in a letter block for syn01 and her matched control, the letters E, N, V and A were presented six times each for 1 s, with 1 s +/- 333 ms fixation only between each letter. This protocol allowed an estimation of the BOLD response to each letter in each block (beta weights, using a General Linear Model, see below) based on six presentations. We obtained three estimations (betas) in each run for each “colour”, for a total of thirty-six estimates (9 * 4 “colours”) for each type of stimulus to be used by classifiers. The power of classification algorithms depends on both the number and quality (signal to noise ratio) of estimates (called exemplars). The present compromise between quantity and quality was based on (Mumford, Turner, Ashby, & Poldrack, 2012) and on preliminary experiments (Ruiz, Hupé, & Dojat, 2012). Subjects had to fixate the centre of the screen (the fixation point, present between stimuli and at the centre of the coloured rings, or the centre of the grapheme) and pay attention to the stimuli for the whole duration of each run. To help subjects maintain attention, they performed a one-back task (pressing a button each time the same stimulus was repeated twice in a row).

In the remaining session, a high-resolution, high-contrast structural image of the brain was acquired using a T1-weighted MP-RAGE sequence. The sequence parameters were TR/TE/TI: 25/3.7/800 ms, excitation pulse angle: 15°, acquisition matrix: 180 sagittal slices of 256*240 (read x phase), bandwidth: 191 Hz/pixel, readout in anterio-posterior direction, number of averages: 1, sense factor anterio-posterior: 2.2, right-left: 2, isotropic nominal resolution: 1 mm, with a total measurement time of 9 min 41 s. This image was the structural reference image of each subject. We also acquired diffusion-weighted images, analysed in another study (Dojat, Pizzagalli, & Hupé, 2018) and a sequence of functional resting state (not analysed yet).

We recorded oculomotor signals during the scans with an ASL EyeTracker 6000. At the beginning of each session, subjects had to fixate each point of a calibration matrix, and were therefore aware that the quality of their fixation was monitored. However, signal quality in some subjects was not good enough or not constant, or even too poor to be of any use for subjects who had to wear non-magnetic glasses in the scanner, so we did not even attempt to analyse these data. We can only speculate that subjects had a better fixation than if they did not know that their gaze was recorded. Whole brain univariate analyses did not reveal any activation along the anterior calcarine and the parieto-occipital sulcus, where activations correspond to the signature of blinks (Hupé, Bordier, & Dojat, 2012a), providing indirect evidence that the distributions of blinks were not correlated with our stimuli presented randomly.

### Data Analysis

The standard pre-processing procedure of functional images was applied using SPM8: slice-timing correction, then motion correction with realignment, together with correction of spatial distortions of the static magnetic field (Vasseur et al., 2010). The within session structural image was realigned to the mean EPI image, as well as the high resolution high contrast structural image, but no further transformation of the EPI images was performed. No spatial smoothing was applied for MVPA, as maximally differential activation of voxels was shown to maximize the power of classifiers (Ruiz et al., 2012). This was confirmed on these data when testing spatial filters with FWHM = 3, 6 and 9 mm. Transformation matrices were computed between the structural image and the MNI template to allow the transformation and projection of atlas-based masks of specific anatomical structures (Anatomy Toolbox for SPM8 Version 2.2b, 2016) into the subject’s space.

For MVPA (our main analysis), for each subject and each run we first ran a General Linear Model (GLM). The six parameters of motion correction were included as factors of non-interest in the design matrix. Thirteen main predictors, four events (grapheme or colour) * three blocks plus one for when only the fixation point was shown, were obtained by convolving the canonical HRF with Dirac functions corresponding to the time of presentation of each stimulus. The corresponding beta weights estimated by the GLM for each colour (real or synaesthetic) and stimulus type (ring or grapheme), divided by the square root of residuals, were used as examples by a Support Vector Classification (SVC) algorithm (Scikit-learn version 0.15.2, implemented in Python version 2.7.9.0: Pedregosa et al., 2011). We used a linear kernel (default value of the C parameter = 1) and a one-versus-one classification heuristic to classify each example in one of the four categories. For all five classifications describes below, training and test runs were always fully independent: betas obtained from blocks from the same run were never split between training and test runs.

#### Classifications

We trained and tested five families of classifiers (Figure 3). Six runs (eighteen blocks) were used for colour (‘Col’ family of classifiers) and synaesthesia (‘Syn’) decoding. The procedure was leave-one-run-out. Six classifiers were therefore trained to classify (5 runs * 3 blocks * 4 colours = 60) colour exemplars in four categories, and tested on (1 run * 3 blocks * 4 colours = 12) independent exemplars. Performance was therefore computed over seventy-two classifications (6 classifiers * 12 tested exemplars), with chance level = 25% and 95% Confidence Interval of chance for each subject = [16 36]% (binomial probability, Agresti-Coull estimation). For grapheme runs, training was performed on pairs made of one letter and one digit. If the decoder learnt only the letters, for example (by being able to filter out the responses to digits), then performance on decoding letters and digits could reach up to 50%, without knowing anything about synaesthetic colours. One could expect, however, that performance of synaesthetes would be higher than for controls because of the additional information provided by synaesthetic colours. A more stringent test of synaesthetic coding (‘g1g2’) was the training of one classifier on letters (3 runs * 12 exemplars) and testing on digits (and the reverse). Learning was achieved using thirty-six exemplars (letters or digits) to be classified in four categories, test was on thirty-six exemplars (digits or letters), for a total performance over seventy-two classifications by combining training on letters and training on digits. To evaluate if brain regions coded both real and synaesthetic colours (‘C2S’), training was performed by one classifier on six colour runs (seventy-two exemplars), test on six grapheme runs (seventy-two exemplars). We also performed the reverse classification (‘S2C’).

**Figure 3.**
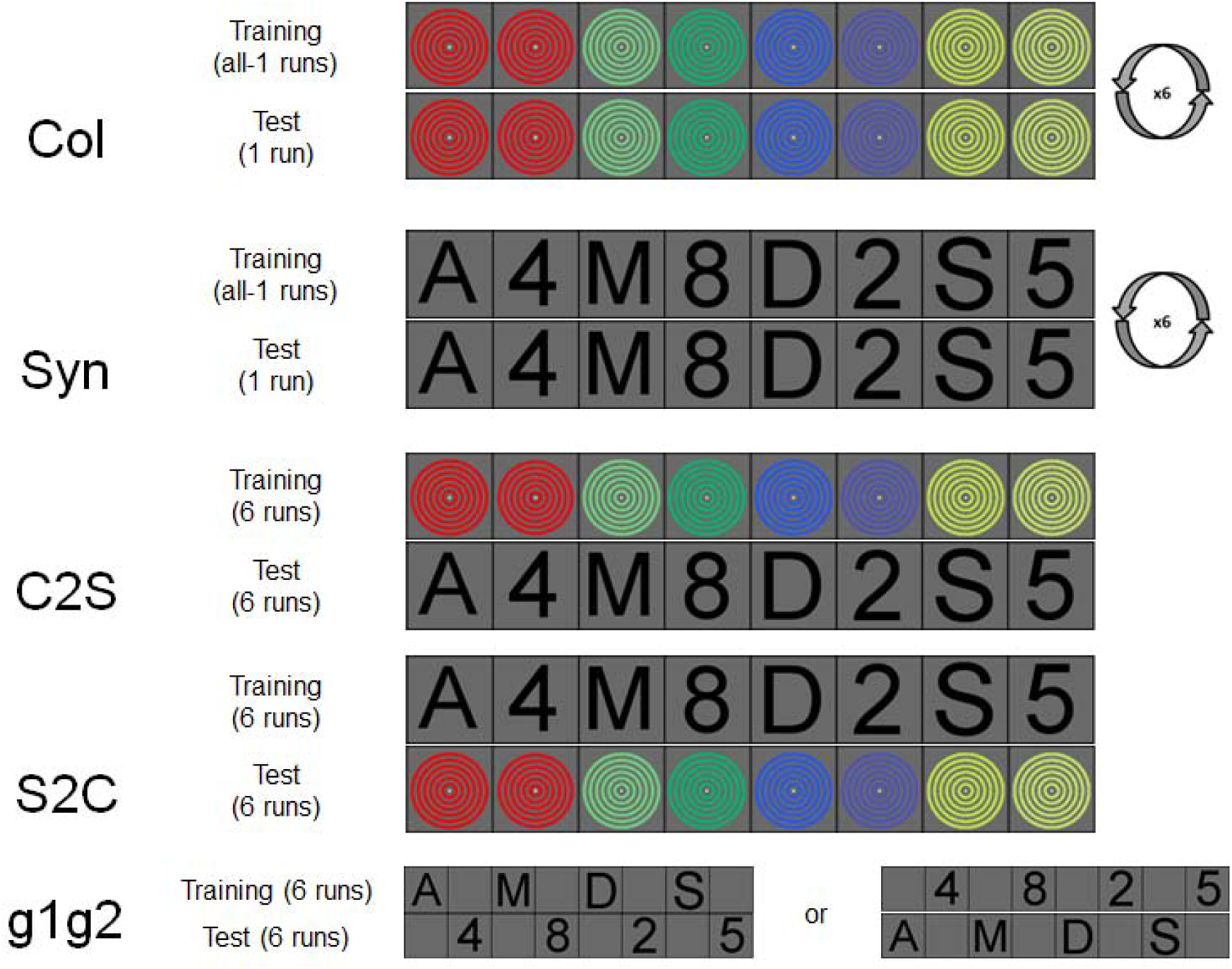
MVPA classifications. ‘Col’ classification: The procedure was leave-one-run-out. Six classifiers were trained to classify 60 colour exemplars from 5 runs in four categories, and tested on 12 independent exemplars of the remaining run. Performance was therefore averaged over seventy-two classifications (6 classifiers * 12 tested exemplars). ‘Syn’ classification: The procedure was the same as for the ‘Col’ classification, based on pairs of graphemes and therefore also synaesthetic colours for synaesthetes. ‘C2S’ classification: Training was performed by one classifier on six colour runs (seventy-two exemplars), test on six grapheme runs (seventy-two classifications). ‘S2C’ classification: The procedure was the same as for the ‘C2S’ classification. ‘g1g2’ classification: Classifiers were trained on letters (3 runs * 12 exemplars) or digits (3 runs) and tested respectively on digits or letters. Overall performance was based on seventy-two classifications.

We computed MVPA in regions of interest (ROIs) defined in each native (non-transformed) subject space. We used visual areas defined by individual retinotopic mapping as well as atlas-based ROIs (Figure 4). We expected synaesthetic colours to involve the ventral visual pathway, anterior to V4, so we tested the four subdivisions of the fusiform gyrus (FG, Figure 4a). Some studies have also suggested the role of parietal areas, even though no consensus emerged about exactly which part if any may be involved (Hupé & Dojat, 2015), so we defined ROIs in parietal regions (Figure 4b).

**Figure 4.**
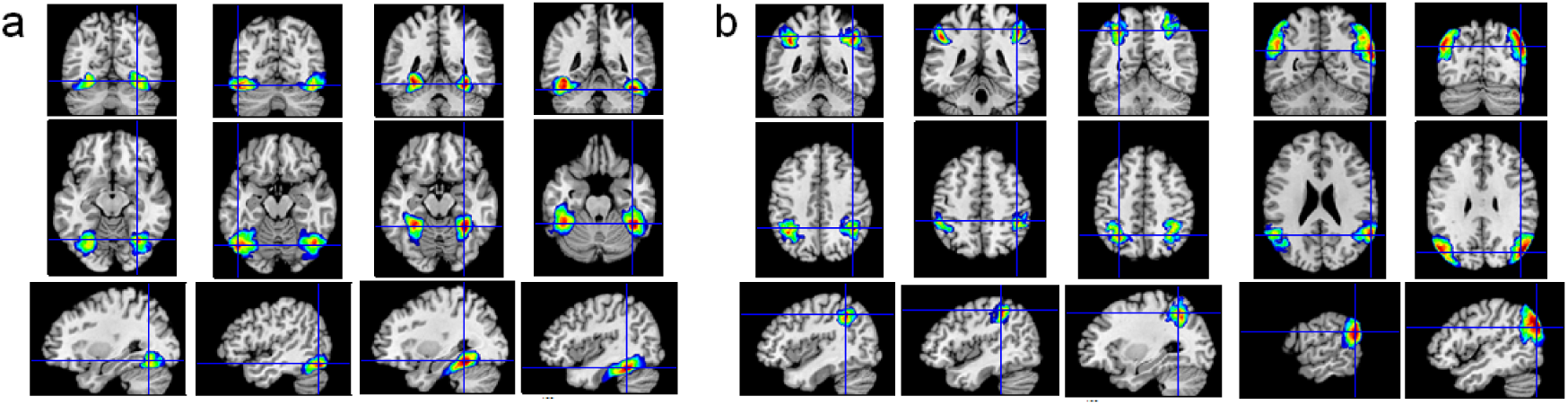
**a.** Atlas-based regions of interest (ROI) of the fusiform region. From left to right, FG1, FG2, FG3 and FG4. Colour gradients denote the probability of being in the specified ROI, from 0% (dark blue) to 100% (dark red). We considered the largest ROI as the mask of the corresponding region. **b.** Parietal ROIs. From left to right, AIPS_IP1, AIPS_IP2, AIPS_IP3, IPL_PGa and IPL_PGp. See text for full names and references of these areas.

For each subject, anatomical ROIs were defined as the intersection of the subject’s grey matter mask and the mask of the anatomical ROIs (Anatomy Toolbox for SPM8: Eickhoff et al., 2007) projected into the subject’s space. Both retinotopic and atlas-based ROIs had different number of voxels within and across subjects. The performance of classifiers may depend on the number of voxels (called “features” for the algorithm), making difficult the comparison of absolute performance in different ROIs. Between-subject differences may also bias group comparisons.

To address this issue, we first tested ROIs of different sizes by regrouping retinotopic areas and subdivisions of the fusiform areas and of the parietal areas. The pattern of results were similar whatever our grouping choice of ROIs. We present the results for ROIs of intermediate size (we indicate the min and max number of voxels across subjects in each ROI), regrouping the right and left parts of retinotopic areas (V1 = [206 441], V2 = [125 420], V3 = [156 340], V4 = [94 268]), the two posterior (J. Caspers et al., 2013) (left = [98 150], right = [61 125]) and anterior (Lorenz et al., 2017) (left = [174 331], right = [127 271]) parts of the fusiform areas, the 3 subdivisions of the Intaparietal Sulcus (Choi et al., 2006; Scheperjans et al., 2008) (left = [197 311], right = [191 275]) and the anterior and posterior parts of the Inferior Parietal Lobule (S. Caspers et al., 2006; S. Caspers et al., 2008) (left = [131 335], right = [124 279]).

We also defined ROIs using the same number of voxels in each subject and ROI. To do that, for all classifications, we selected 100 voxels with the highest F-scores to colours in each area (we tested different selection sizes and found that 100 was about the optimal number of voxels to reach maximum performance). In order to have enough voxels to choose from in every subject, we selected voxels in only six large areas: the left and right retinotopic areas V1 to V4 (minimum number of voxels across subjects were respectively 352 and 327), the left and right fusiform areas FG1 to FG4 (298 and 188) and the left and right parietal areas (347 and 315). Such a selection provides the best chances for colour classifiers (since we select voxels maximally modulated by colours), but classification is then not independent of selection when measuring colour decoding after selection of F-scores to colours (but classification is independent for grapheme decoding). In order to provide a fair measure of colour decoding performance to compare grapheme decoding with, voxels were selected using F-values computed based only on runs used for training, meaning that each of the six training sets was based on a different set of voxels. For other classifications, the same set of voxels was used based on F-values computed across all colour runs.

### Statistical analysis

In each ROI, we computed 95% CIs of the performance of each group, as well at the 95% CIs of the between group differences. We performed both independent and paired comparisons. Paired comparisons are in principle more appropriate and powerful with this protocol, because it cancels any difference due to the specific choice of colours and graphemes; however, for voxels not concerned with those small differences, pairing is artificial and may just bring some noise. Results were in any case very similar for both comparisons. We show the CIs for paired comparisons. We also performed paired comparisons by computing the 95% CI of the odds ratio when comparing 19 synaesthetes against their matched controls, using a mixed-effect generalized linear model, with a binomial family and a logit link function, as implemented in the library lme4 (Bates, Mächler, Bolker, & Walker, 2015) in R, version 3.3.3.

In order to fully exploit our data set, we performed two additional analyses.

A searchlight analysis was performed over the whole brain (Kriegeskorte et al., 2006). Whole brain analyses are in principle less powerful than ROI analyses because they constrain to distort each subject’s anatomical space within one common space, so the average performance at any given voxel may in reality correspond to different anatomico-functional voxels in different subjects. Moreover, they re-introduce the methodological issues related to spatial smoothness (Stelzer, Lohmann, Mueller, Buschmann, & Turner, 2014). This analysis was therefore exploratory. It allowed us to discover other clusters potentially involved in synaesthesia, which we could further analyse as post-hoc regions of interest to see if they displayed a consistent pattern of results across classifiers. The searchlight analysis used a 15 mm radius and the SVC algorithm. Performance maps were transformed to the common DARTEL space for voxel-wise group comparisons (resolution 3 by 3 by 3 mm). We performed in SPM8 two-sample (groups of 20 subjects) and paired-sample t-tests (N = 19) between synaesthetes and controls, as well as one-sample t-tests to compare the average performance of each group (N = 20) against chance (= 0.25). For all comparisons, no individual voxel reached pFWE < 0.05. We used cluster-based statistics with the cluster-forming threshold set to p = 0.001 and pFWE < 0.05.

We also performed whole-brain univariate analyses on the groups of synaesthetes and controls to test for differences of magnitude of the BOLD responses to graphemes evoking synaesthetic colours. The design of the experiment was not optimized for these analyses since we did not have any control stimuli (those being not necessary for MVPA). The rationale was the same as for the whole brain searchlight analysis: if any difference was found between synaesthetes and controls, the revealed clusters could be defined as post-hoc regions of interest for our classifiers to test if those regions were involved in coding synaesthetic colours. A 9 mm FWHM spatial smoothing was applied to the subjects’ EPI images before testing two contrasts: a T-contrast of all stimuli against the fixation point (we did not have graphemes that did not evoke any synaesthetic colour); an F-contrast of the four pairs of graphemes. Contrast maps were distorted within the study-specific template computed using DARTEL procedure as implemented in VBM8 (Dojat et al., 2018) and to the MNI space (resolution 1.5 by 1.5 by 1.5 mm). For second-level analyses, we compared the contrast maps of synaesthetes (N = 20) against controls (N = 20) using t-tests (testing stronger signals either in synaesthetes or controls). We also performed paired t-tests on 19 synaesthetes against their matched control to account for possible differences due to the specific choices of graphemes in each synaesthete. For all comparisons, no individual voxel reached p < 0.05, corrected for the family-wise error (FWE, based on the random field theory as implemented in SPM8). We used cluster-based statistics with the cluster-forming threshold set to p = 0.001 (Eklund, Nichols, & Knutsson, 2016) and pFWE < 0.05. As a final control analysis, we performed the same analyses for coloured stimuli.

#### Data Availability

The datasets generated and analysed during the current study are freely available on request (https://shanoir.irisa.fr/Shanoir/login.seam), contact M. Dojat. The data are not publicly available due to privacy restrictions.

## Results

### Multivariate pattern analysis in regions of interest (defined at the individual level)

Figure 5 shows the performance of all classifiers (described in Figure 3) in all our ROIs, without any voxel selection (ROIs have therefore different number of voxels across regions and subjects).

**Figure 5.**
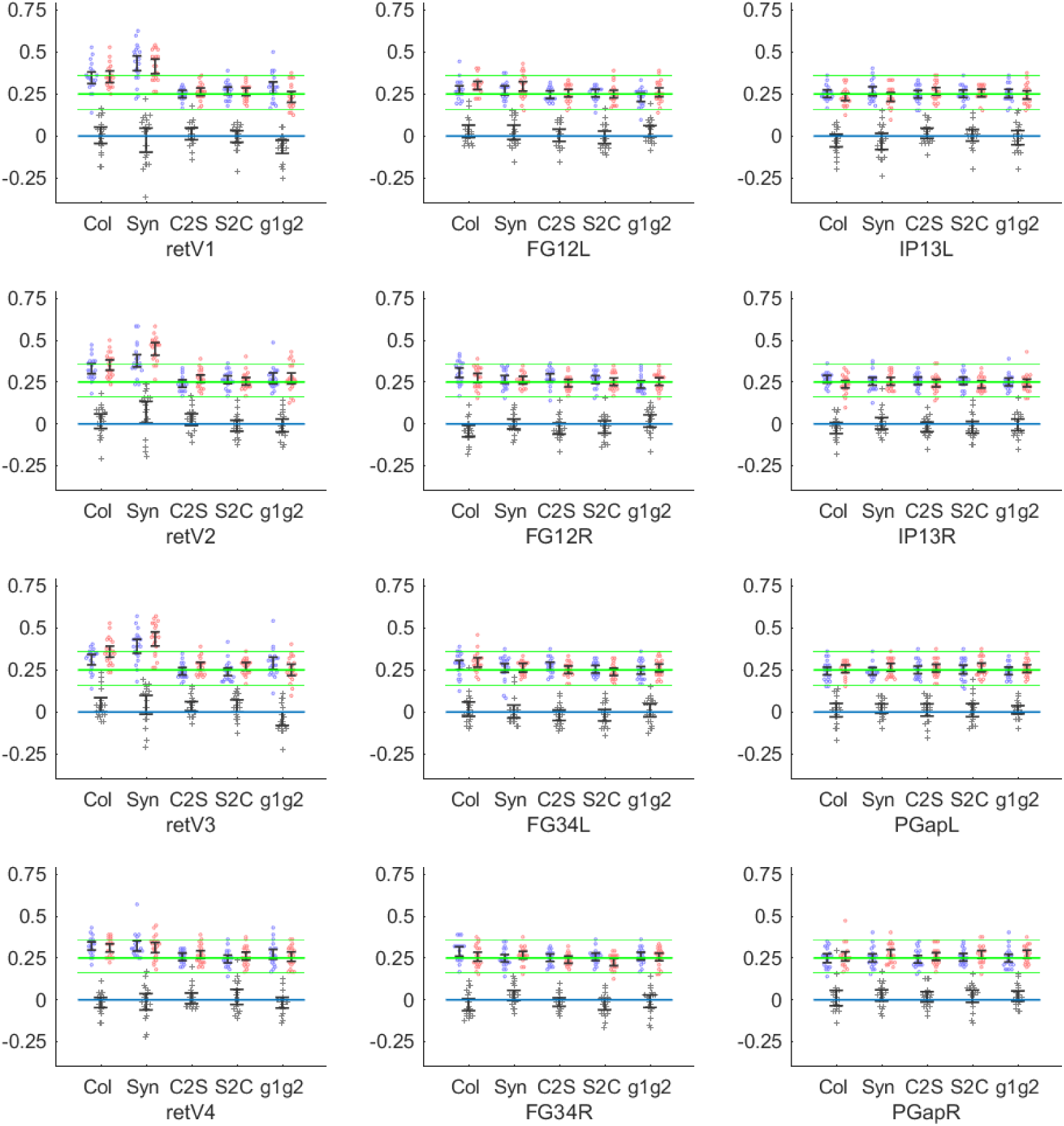
Performance of classifiers in retinotopic areas, the fusiform gyrus and parietal areas. Each ROI regrouped several areas, for example the left and right parts of V1 for ‘retV1’ in order to provide a large number of voxels in each subject and ROI (at least > 60, and > 100 voxels in most ROIs; see Methods: Data Analysis). ‘retV1’ to ‘retV4’ were defined based on retinotopic mapping in each subject; other ROIs were defined as the intersection of the subject’s grey matter mask and the mask of atlas-based anatomical ROIs (Anatomy Toolbox for SPM8) projected into the subject’s space (see Figure 4). FG12L = left (FG1 + FG2), IP13L = left (AIPS_IP1 + AIPS_IP2+ AIPS_IP3), PGapL = left (IPL_PGa + IPL_PGp), etc. Each classifier was trained and tested on beta weights computed on voxels in the native subject space with no spatial smoothing. Each panel displays the individual and average performances of five classifiers (see Figure 3): ‘Col’ = training and test on betas for real colours (rings); ‘Syn’ = training and test for synaesthetic colours (graphemes, letters or digits); ‘C2S’ = training on real colours (rings) test on synaesthetic colours (graphemes); ‘S2C’ = training on synaesthetic colours test on real colours; ‘g1g2’ training on letters test on digits or training on digits test on letters. The y-axis represents both the *performance* of classifiers (between 0 and 1, chance level = 0.25, thick green line; 95%CI of chance for each subject = [0.16 0.36], thin green lines) for individual subjects (blue = controls, red = synaesthetes) and their group average (with 95% Confidence Intervals) and the *difference of performance* (grey crosses) between synaesthetes and their matched controls (0 = no difference between groups, blue line; whiskers denote 95% CI).

In each subplot, the first two beeswarms from the left show the performance of decoders in each subject for real colours (‘Col’ classification). As expected, the decoding of real colours was above chance (0.25, thick green line) in retinotopic areas as well as in the fusiform gyrus for both controls (blue points) and synaesthetes (red points). No difference was expected nor observed between groups (whiskers across the zero blue line denote the 95% CI for paired comparisons of performance of 19 synaesthetes against their matched control, the difference of performance being denoted by the grey crosses). Note though that the whole 95% CI was slightly above 0 in retinotopic V3, ‘retV3’, and it was slightly below 0 in the subdivisions 1 and 2 of the right fusiform gyrus, ‘FG12R’ (differences are more visible when estimating the CI by a mixed-effect generalized linear models: Supporting Information, Figure S4). But without any independent evidence, these small differences could be due to random sampling. Indeed, all the 99.58% CIs included 0 (Bonferroni correction over 12 tests).

The next beeswarms are for the classification of pairs of graphemes. In synaesthetes only, classification could in principle be achieved based on the synesthetic colours, since the synesthetic colour was the main shared feature associated to each grapheme pair (in most cases one letter and one digit) had in common (so we called it the ‘Syn’ classification). For example, E and 7 were both associated to red by syn01 (see Figure 1). Performance was above chance level (25%) in controls. This means that this performance could be achieved by classifiers based on either some spatial features shared by each grapheme pair or by optimizing decoding to only one of the graphemes. In order to test whether graphemes could be decoded on the basis of synaesthetic colours, we looked if synaesthetes performed better than controls. This was the case in retinotopic V2 (95% CI of the difference of performance, two-sample t-test: [1.5 12.5]%; paired t-test: [0.5 13.4]%; 95% CI of the odds ratio = [1.15 1.56]; see Methods, statistical analysis) and to a lesser extent in retinotopic V3 (but note that performance was lower for synaesthetes in the subdivisions 1 to 3 of the left Intra-Parietal Sulcus, IP13L; such a difference is most likely due to random sampling since none of the group performances in IP13L was above chance). Only the difference in V2 survived Bonferroni correction over 12 tests, for the mixed-effect analysis (*p* = 0.0002).

The third group of beeswarms represent the data answering our main question: can synaesthetic colours be decoded based on real colours (‘C2S’ classification)? This was not the case in any ROIs we considered in controls, as expected, but also in synaesthetes (the 95% CI of all groups crossed the 0.25 chance baseline). In particular, performance was not significantly above chance in V2, as would have yet been expected if the higher performance in synaesthetes for the ‘Syn’ classification was really due to the coding of synaesthetic colours. Moreover, there was no evidence of better classification in synaesthetes than controls. We obtained similar null results when we tried to decode real colours based on graphemes (and possibly synaesthetic colours in synaesthetes: ‘S2C’ classification).

The last beeswarms on the right of each subplot show the performance for another classification that should have been possible only on the basis of synaesthetic colours: classifiers were trained on one set of graphemes (digits or letters) and tested on the other set of graphemes (‘g1g2’ classification). Again, performance was never above chance and we found no difference between groups, in particular in V2. We even observed lower scores for synaesthetes in V1 (where it even survived Bonferroni correction for the mixed-effect analysis: *p* = 0.0002) and V3, where we had yet observed higher performance for ‘Syn’ decoding. This lack of consistency across different tests addressing the same question confirmed that these small variations, even when statistically “significant”, were most likely due to random sampling.

We performed again all these analyses using six larger ROIs (regrouping either the left or the right parts of V1 to V4, FG1 to FG2 and the areas of the inferior parietal lobule and the intraparietal sulcus) in which we selected the 100 voxels with the largest scores to F-tests to real colours, in order to feed the classifiers with the voxels most sensitive to real colours (Figure 6). The performance of the ‘C2S’, ‘S2C’ and ‘g1g2’ classifiers was never above chance in synaesthetes (nor in controls, as expected), and performance was never better in synaesthetes.

**Figure 6.**
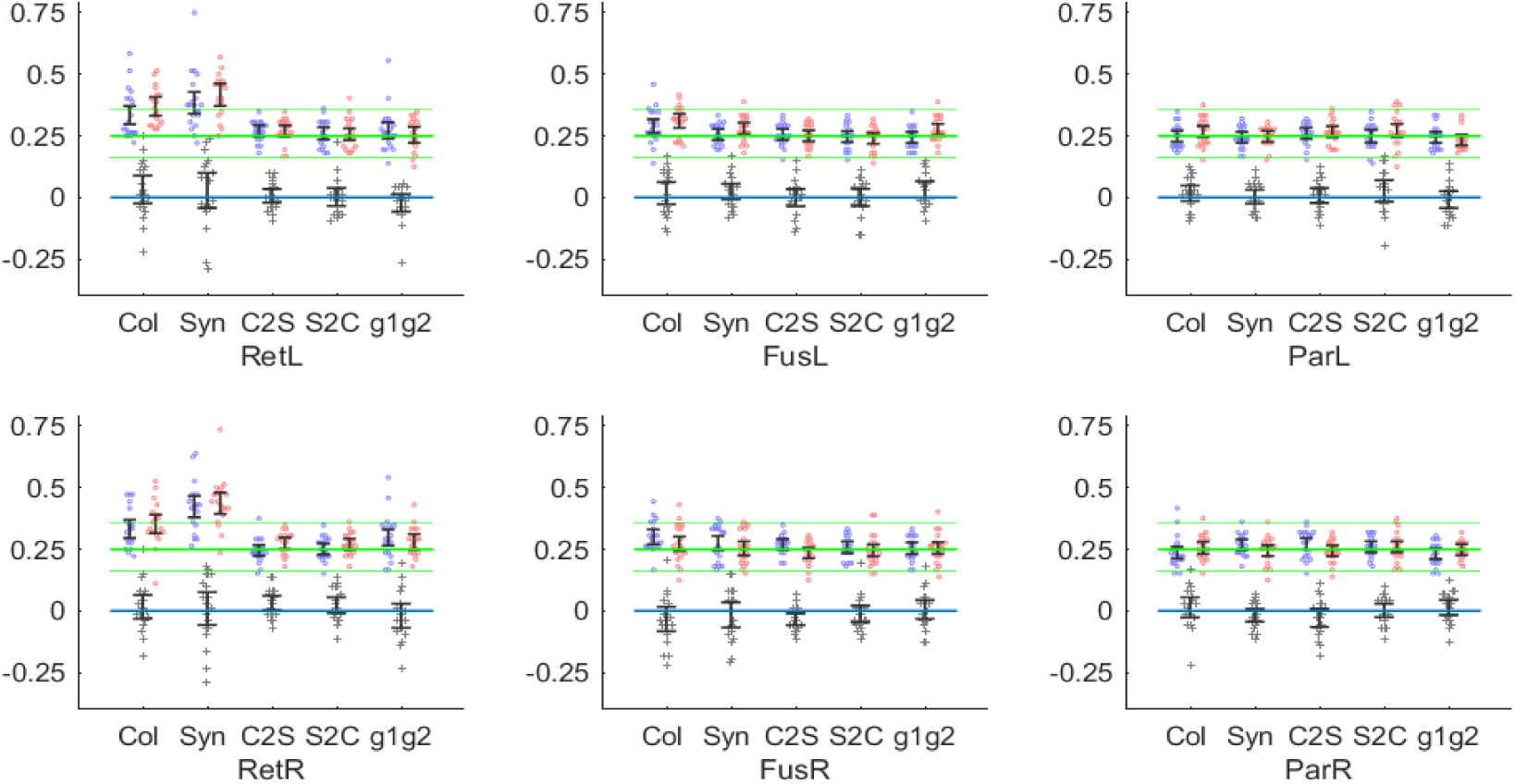
Performance based on the same number of voxels (= 100) in each large ROI (retinotopic areas, fusiform gyrus and parietal regions) and subject. For the classification of real colours (‘Col’), the selection of the best F-values to colours was different for each of the six leave-one-out classifications, based each time only on the five runs used for training the classifier, to insure independence of training and test. For the other selections, all colour runs were used to select the voxels with the highest F-scores. The high performance for the ‘Syn’ classification in retinotopic areas indicates that many voxels respond both to change of colour or luminance and the shape of graphemes, probably thanks to the small receptive fields of lower visual areas. Same conventions as in Figure 5. The CIs of the odds ratio computed by mixed-effect generalized linear models are shown in Supporting Information, Figure S5.

### Individual differences

The phenomenological experience of synaesthetic colours may vary a lot across synaesthetes, which may compromise the visibility of effects at the group level. While this phenomenology has been so far problematic to capture with objective measures, we could estimate the strength of the synaesthetic associations in each subject, using variants of Stroop tasks. We reasoned that synaesthetes with stronger synaesthetic associations might have stronger modulations of the BOLD signal and thus larger decoding values. We first tested area V2, where we had observed on average higher performance in synaesthetes for the ‘Syn’ classification. We were wondering if this higher performance was really due to the coding of synaesthetic colours, because our other, more specific classifiers (‘C2S’, ‘S2’C and ‘g1g2’), had not revealed any difference. A correlation between synaesthetic strength and performance would constitute an independent validation of coding. Figure 7 shows the performance of each subject as a function of the strength of synaesthetic associations, measured in Stroop-like psychophysics experiments (see Table 1 and Ruiz & Hupé, 2015, for further explanations about the ‘Photism Strength’ index) for synaesthetes only (red crosses). Controls (blue circles) were attributed the value of their matched synaesthete.

**Figure 7.**
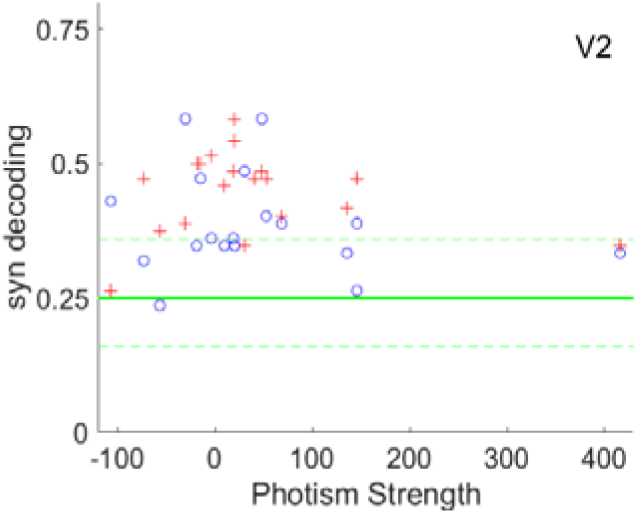
Performance of the classifier trained and tested with synaesthetic colours (pairs of graphemes) in each subject in area V2 defined retinotopically (same data as ‘Syn’ in the second panel of the first column of Figure 5) as a function of the strength of synaesthetic associations (‘Photism Strength’). This strength, measured for synaesthetes (red crosses), does not show any evidence of correlation with the performance of the ‘Syn’ decoder (*r* = −0.11, 95% CI = [−0.54 0.36]). Controls (blue circles) were attributed the value of their matched synaesthete (*r* = −0.18, 95% CI = [−0.58 0.30]). Note that one value of Photism Strength was larger than the other ones. We carefully checked that this value was correct. However, given its possible influence on the correlation results, we complemented this analysis with two other analyses, by removing this value (for synaesthetes, *r* = 0.28, 95% CI = [−0.21 0.66]) and by performing non-parametric correlations (Spearman *r* = −0.03, N = 19, *p* = 0.90). We compared the results of these three statistical tests for all the other tested correlations. The statistical conclusions were always similar, except for one case described in Supporting Information, Figure S2.

There was no correlation between both measures, neither for synaesthetes nor, as expected, for controls. We also observed that the difference of score between each synaesthete and her (or his) matched control did not increase with photism strength (*r* = 0.06, 95% CI = [-0.42 0.52]; Spearman *r* = 0.01, N = 18, *p* = 0.95). Therefore, this analysis did not provide any independent argument in favour of the decoding of synaesthetic colours in V2. We computed similar correlation analyses in every ROI and for all classifiers and never found any correlation (all uncorrected *p* > 0.05). We also computed both positive and negative correlations over the whole brain for the five classifiers, independently for synaesthetes and controls. We never found any significant cluster (cluster forming threshold, *p* = 0.001).

### Whole brain searchlight multivariate pattern analysis

We complemented our ROI analysis with searchlight analyses over the whole brain (normalized to the MNI space), comparing the performance of each group against chance as well as comparing groups for the five classifications. These exploratory analyses were performed to discover clusters potentially involved in synaesthesia outside of our ROIs. “Significant” clusters could be used as post-hoc regions of interest to see if they displayed a consistent pattern of results across classifiers (these results are displayed in Supporting Information, Table S1). We found no differences between controls and synaesthetes at our statistical threshold for classifiers trained and tested on colours (rings, ‘Col’ classifiers). For classifiers trained and tested on synaesthetic colours (graphemes, ‘Syn’ classifiers), we observed higher performance in synaesthetes in the parietal cortex (Table S1: on the right side with paired t-tests and on the left side with two-sample t-tests; bilateral difference could be observed for both contrasts when using a higher cluster-forming threshold). However, testing synaesthetes against chance revealed no cluster at our threshold around these coordinates of the parietal cortex (performance was above chance in both groups in the occipital cortex, as expected).

We found no difference between controls and synaesthetes at our statistical threshold for the critical test of shared coding of real and synaesthetic colours, when classifiers were trained on coloured rings and tested on graphemes (‘C2S’ classifiers). Testing synaesthetes against chance also revealed no cluster. The reverse classification (learning on graphemes, ‘S2C’), however, revealed two clusters with higher performance in synaesthetes for independent t-tests, in the right occipito-temporal cortex and in the left putamen. Only the first cluster was confirmed by paired-comparisons. When testing performance against chance two clusters emerged for synaesthetes (none for controls), one again in the same part of the right occipito-temporal cortex, and the other in the left parietal cortex, abutting the parietal cluster obtained previously for the higher performance in synaesthetes for the ‘Syn’ classification (we shall come back to this concordance in the following *post-hoc* analysis).

Finally, for classifiers trained on either letters or digits (and tested respectively on either digits or letters), a critical test of the coding of synaesthetic colours, higher performance was observed, but in controls, in the left inferior frontal gyrus, for both paired and independent t-tests. However, no cluster emerged anywhere in the brain in controls (nor in synaesthetes) when testing performance against chance, so this cluster should be considered as a false positive.

*Post-hoc* analysis. We further explored the performance of classifiers in the two clusters identified by the ‘Syn’ classifier and the five clusters identified by the ‘S2C’ classifier, corresponding in fact to two parietal regions (left and right), one right occipito-temporal region and one cluster in the left putamen. In each cluster, we computed the average across voxels of the searchlight scores, in order to compare the performances of our five classifiers for synaesthetes and controls in these seven clusters defined *post-hoc*, with two-sample and paired-sample t-tests. We also compared the performance of each group against chance. Statistically “significant” differences were obtained only for the contrasts used to define the clusters (Table S1). Only one additional comparison was “significant” (*p* = 0.012, not corrected for multiple comparisons) in the left parietal cluster at XYZ = [−33 −28 50], which had been obtained when testing synaesthetes against chance for the ‘S2C’ classification (training classifiers on graphemes and testing them on colours: Figure 8): synaesthetes also performed better than controls at decoding graphemes (‘Syn’ classification), 95%CI = [1 9]% (paired comparisons), and better than chance (95% CI = [26 31]%), but the performance was not correlated with the strength of synaesthetic associations (*p* = 0.51).

**Figure 8.**
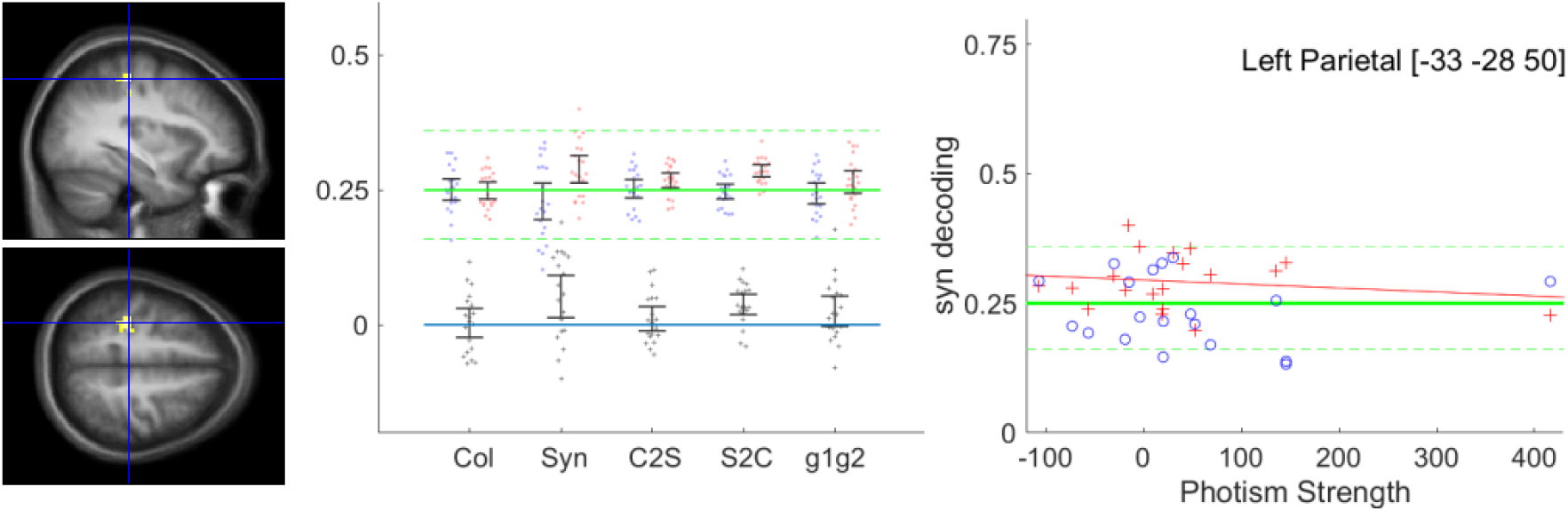
**Left**: Parietal cluster identified based on whole brain searchlight analysis for ‘S2C’ decoding, Synaesthetes>chance (27-voxel cluster at XYZ = [−33 −28 50], one-sample t-test). **Middle**: performance of classifiers in this cluster (same conventions as in Figure 5). The performance of synaesthetes was logically above chance for the ‘S2C’ classification, since the cluster was defined based on this contrast. For the independent classifier ‘Syn’, the performance of synaesthetes was also above chance and above that of controls. **Right**: Absence of correlation between the strength of synaesthetic associations and ‘Syn’ decoding (Spearman *r* = 0.02, N = 19, *p* = 0.95; for ‘S2C’ decoding, not shown: Spearman *r* = −0.15, N = 19, *p* = 0.53).

### Whole brain univariate analyses (normalized anatomical space)

Similarly to the searchlight analyses, these whole-brain univariate exploratory analyses were performed to discover clusters potentially involved in synaesthesia outside of our ROIs. They revealed no difference between controls and synaesthetes at our statistical threshold for T-contrasts of graphemes when performing two-sample t-tests. However, the paired-sample t-tests revealed stronger BOLD signal in a small cluster in synaesthetes, close to the left precentral gyrus. We treated this cluster as a candidate region for the coding of synaesthetic colours (Supporting Information, Table S1).

For F-contrasts, we did not observe any stronger modulation in synaesthetes (neither for two-sample nor paired-sample t-tests). Surprisingly, we observed stronger modulation in controls in two clusters (paired comparisons), in the right occipito-parietal cortex (Supporting Information, Figure S1) and in the left insula. The two-sample t-tests revealed only the occipito-parietal cluster. We did not have any explanation for these differences, which might be false-positives (Eklund et al., 2016). We note that the analysis by (Rouw & Scholte, 2010) revealed a cluster (which they called IPS, cluster extent = 3280 mm3) at equivalent peak coordinates on the left side ([−30 −72 28]), obtained with the contrast synaesthetes>controls for (synaesthetic graphemes)>(non-synaesthetic graphemes). In our case, the weaker modulation by graphemes in synaesthetes would rather argue against the hypothesis of a functional role of this region in synaesthesia. We included these two regions in our post-hoc MVPA analyses for further exploration.

We also tested T- and F-contrasts for the responses to real colours (rings). We observed stronger BOLD signal (T-contrast) in synaesthetes only, in three clusters for paired comparisons (in the left posterior and anterior insula – see Supporting Information, Figure S2 - and in the left parahippocampal region) and two other clusters for two-sample t-tests (in the right middle temporal gyrus and in the right superior, medial, frontal gyrus - see Supporting Information, Figure S3). The lack of consistency between paired and two-sample t-tests could again suggest false-positives, but we nonetheless included these five clusters in our post-hoc MVPA analyses, in case those stronger activations be related to the implicit activation of graphemes by the colours associated to them (“bi-directional” synaesthesia: Gebuis, Nijboer, & Van der Smagt, 2009). F-contrasts to colours revealed only one cluster of stronger modulation in controls in the frontal region, but in the middle of white matter and thus clearly a false positive.

We compared the performances of synaesthetes and controls for our classifiers in those eight clusters defined post-hoc, with two-sample and paired-sample t-tests. Only three comparisons came out “significant” at p < 0.05, but without correction for multiple comparisons (Table S1). Supporting Information, Figs. S1 to S3 detail the results obtained in these three post-hoc clusters.

## Discussion

Our goal with this study was to provide univocal evidence of the activation of the ‘colour network’ by imaginary colours as experienced by grapheme colour synesthetes.

Studies based on univariate analyses to search the neural correlates of synaesthetic colours face two major problems. First, BOLD responses to stimuli leading to the experience of synaesthetic colours need to be compared to a control response (subtraction method: see the Figure 1 by Hupé & Dojat, 2015). Such a control response may be obtained by testing the same subjects with similar stimuli that do not generate a synaesthetic experience (pseudo-graphemes or graphemes that, by chance, do not generate such an experience in the tested synaesthetes). The problem is, it is impossible to know whether the additional activations, if observed, are specific to the synaesthetic experience of colours. For example, letters and numbers can also be named, unlike pseudo-graphemes. A control response may also be obtained by testing non-synesthetes with the same stimuli. But, again, it is impossible to know whether the additional activations, if observed, are specific to the synaesthetic experience of colours. For example, synaesthetes often enjoy visualizing the synesthetic colours of graphemes: attentional and emotional components may therefore bias the comparison. Second, averaging the results across subjects require to transform the individual data within a common reference space, with the possible loss of fine-grained spatial information. The first problem may generate false positive results, the second false negative results.

Thus, because univariate analysis led to inconsistent results (Hupé & Dojat, 2015), we used in this study Multivariate Pattern Analysis (MVPA) on 20 synaesthetes and 20 control participants, to explore whether the neural processing of real colours and synaesthetic colours shared patterns of activations. To our knowledge, it was the first time that MVPA was proposed in this context. By using MVPA, we could in principle overcome problems associated with univariate analysis because we test the performance, in each individual, related to the coding of certain attributes, not a level of activation in need of further interpretation. Our design was optimized to test if classifiers, trained to distinguish patterns of responses to four different real colours in groups of voxels from different regions of the brain, could classify above chance the responses of those voxels to achromatic graphemes leading to the synaesthetic experience of the exact same colours. The logic was that only synaesthetes tested with their exact, idiosyncratic, synaesthetic code could produce above-chance performance. Unfortunately, the performance of our group of 20 synaesthetes remained very close to chance (all p < 0.05, uncorrected) in all our selections of voxels (retinotopic areas defined at the individual level as well as the fusiform gyrus and parietal regions of interest defined based on a probabilistic atlas), and whatever the extent of the chosen areas or the selection method of the voxels (Figure 5 and 6, classification ‘C2S’ in synaesthetes shown by red points).

A statistical comparison revealed that the performance of synaesthetes was also no better on average than that obtained in control subjects. Of course, the absence of statistically significant effect cannot lead to conclude to the absence of effects. The null result could be due to a lack of power, if, for example, only a few synaesthetes had reached a good performance. However, the results were striking when inspecting the distributions of individual performances: the scores of most synaesthetes were distributed around the chance level, and almost no synaesthete reached an individual performance above the chance level (binomial probability: all the points are included within the green dotted lines). The correlation analyses with a measure of individual differences (the strength of the synaesthetic associations) further confirmed the homogenous performance of synaesthetes around chance. Under our experimental conditions, the collected data therefore do show quite convincingly the absence of shared coding of real and synaesthetic colours.

We further analysed our data set in different ways, in order to be able to detect some signs of coding of synaesthetic colours by neural networks not involved in the coding of real colours. The ‘g1g2’ classification, which could have been achieved only on the basis of shared synaesthetic colours across letters and digits, remained at the chance level in synaesthetes. The ‘syn’ classification, expected to reach a higher performance in synaesthetes, was similar in controls. We also explored the performance of classifiers beyond our regions of interest, across the whole (normalized) brain (searchlight analysis) without obtaining significant results.

We also explored the whole brain using mass univariate tests, knowing that whole brain analyses face the ill-posed problem of correction of multiple comparisons of partly correlated tests, problem not fully solved by the Random Field Theory (Eklund et al., 2016). Moreover, since we performed in total at least nine whole brain searchlight analyses and four whole brain univariate comparisons (T- and F-contrasts for responses to graphemes and colours, see Table S1), we could have set a family-wise error level at 0.05/13. We preferred to keep a non-corrected level for easier comparisons with other studies. The whole brain analyses were used only for exploration, and for every detected cluster we searched for additional evidence (differential response for other comparisons, or correlation with individual differences). Since we did not find any additional evidence, we conclude that these clusters may be false positives. We however mention them (see Table S1) in case additional evidence be found in other studies.

For now however, our results further suggest that 3T fMRI studies may not be able yet to identify the neural correlates of the synaesthetic experience of colour (Hupé & Dojat, 2015), probably because those are fine-grained distributed at a resolution lower than our 3-mm voxel resolution, or because the nature of its coding does not translate (well) into BOLD responses. Surprisingly, though, if considering synaesthetic colours simply as a form of mental imagery, we were expecting above-chance decoding performance as observed for other tasks involving mental imagery. Those other tasks, however, typically involved different categories of objects, like food, tools, faces and buildings (Reddy et al., 2010) or objects, scenes, body parts and faces (Cichy, Heinzle, & Haynes, 2012), which evoked stronger BOLD signal in specific areas (like the Fusiform Face Area). Other studies involved retinotopic properties (Thirion et al., 2006) where, again, differences of BOLD signal can be easily observed. Here we were trying to decode mental images within only a single category, colour. This confirms that synaesthetic experiences do not evoke strong BOLD responses, at least when using standard 3T MR scanner, as already suggested by the inconsistency of the published results based on univariate models.

Below we consider alternative explanations (e.g., methodological flaws) to the absence in our data of neural traces of the processing of synaesthetic colours.

### Colour imagery could not be decoded

Our study used a protocol very similar to that used by (Bannert & Bartels, 2013), who could decode the typical colour from eight objects, presented as greyscale photos, with classifiers trained on concentric colour circles designed after (Brouwer & Heeger, 2009), like in our study. The prototypical colour of the objects was red, green, blue or yellow (like a banana and a tennis ball). Across 18 subjects, decoding accuracy was “significantly” above chance in V1, but reached only 32% on average, which is hardly above the 95% CI ([24 30]%) of the performance observed for our similar classifier (‘C2S’) in the areas V1 to V4 of synaesthetes. Their experimental procedures, slightly different from ours, may have better optimized the signal to noise ratio and allowed this higher performance (see below). Alternatively, since the colour-diagnostic objects were presented before the coloured concentric rings, subjects may have imagined, when viewing the rings, the very objects that were presented before. Subjects had to do a motion discrimination task to divert their attention (similarly to our one-back task), but such a task (like ours) was not very demanding (though note that Bannert and Bartels argue that their results are due to automatically occurring processes during object vision rather than active imagery). Of course, a similar argument holds even more in our experiment: synaesthetes were very likely to recognize the colour matching exactly their synaesthetic colour of letters and digits, and they might well have imagined the letter or digit when looking at the coloured stimuli. In both cases, decoding would be based on the complex shape of stimuli rather than their colour. In the case of Bannert and Bartels, objects were similar to those used in other successful visual-to-imagery decoding and involved several categories of objects as well as different retinotopic properties (the objects had different orientations but were rotating; however the banana or the coke can, for example, had about 12 deg extent, apparently much more than the Nivea tin or the blue traffic sign), while in our case objects all belonged to the grapheme category, and all spanned the same visual extent. It is therefore possible that in the study by Bannert and Bartels the slightly above chance decoding performance was due to residual category and retinotopic properties, not to colour. With such an interpretation, decoding of imaginary colours would have failed in both their and our study.

### Flaw in the paradigm used: duration of event presentation

Our close to chance performance could be linked to our choice of a fast event related paradigm, each stimulus being presented each time for only 1 s, with an ISI = 1 s +/- 333 ms. Bannert and Bartels presented images for 2 s with a 1 s ISI, each repeated four times in a row (miniblocks). One may wonder whether our presentation time was sufficient to trigger synaesthetic associations. However, psychophysical tasks show that the naming of the synaesthetic colours of graphemes takes on average much less than 1 s (Table 1). Because of our one-back attentional task, though, we cannot be sure that the synaesthetic associations were always conscious. However, synaesthetes did not report any specific difficulty with their synaesthetic experience when viewing, inside the scanner, the proposed paradigm. We designed such a protocol because we did not want synaesthetes to pay too much attention to their synaesthetic colours, then possibly triggering complex attentional and emotional processes. Those components are part of the synaesthetic experience, but they do not tell us anything about the phenomenological experience of colours, our main goal being to try to isolate the possible neural commonalities of the real and synaesthetic experience. The quasi-absence of observed differences of overall activation and modulation between synaesthetes and controls for graphemes indicates that we were successful in synaesthetes having a similar experience to controls for graphemes, in terms of attentional and emotional content. With different conditions, favouring synaesthetic colours to be experienced intensely, we would expect the overall pattern of brain activity to be different, but those differences would be poorly informative.

### Flaw in the paradigm used: type of stimuli

A critical aspect of our fast-event paradigm is related to the slow dynamics of the hemodynamic signal and the signal to noise we could obtain. Here, the critical benchmark was the possibility to decode real colours, since the protocol was identical for synaesthetic and real colours. We were successful in decoding colours above chance in the visual cortex, but not to the extent that we hoped: only 35% on average, chance being 25%. Using 12 s miniblocks, Bannert and Bartels obtained an average performance for colours between 35% and 40% in V1 to V4. Differences other than the timing of the stimuli may explain this only slightly higher performance: their total presentation time of coloured stimuli was about 42 min (20 min in our study; for example, Brouwer & Heeger, 2009, obtained even higher performances with experienced participants tested for much longer durations); their stimuli were much larger (7.19 deg vs. 2 deg radius) and isoluminant (we do not know whether luminance information in our case helped or hindered decoding). Because they were constrained by the idiosyncratic synaesthetic associations, our stimuli were also not well distributed within the colour space (see Figure 2). Colour differences between categories (R, G, B and Y) and similarity between colours for pairs (letter-digit) were different between subjects and not always optimal to reach maximal performance by classifiers. Probably, some pairs of supposedly similar colours confused classifiers, as well as short distances between some categories. Retrospectively, we should in fact even consider ourselves lucky to have achieved such a performance for colour decoding. Choosing only three colours would have allowed us to avoid confusions and get more exemplars for each colour (with fewer categories to decode, though, confounding factors are more likely). More repetitions would be welcome, however we wanted to record signals for real and synaesthetic colours within the same scanning session to avoid any spatial smoothing of the voxels (which is often necessary when aligning images obtained in different sessions). Preliminary experiments had showed us indeed that combining the signals from different sessions did not improve performance (Ruiz et al., 2012). Our total session time was about 1 hour, which is about the limit one may ask naïve subjects to lie in a scanner without moving while maintaining fixation and attention over boring stimuli.

### Lack of statistical power to detect small differences

Given our moderate performance for colour classification, our absence of above-chance performance for the decoding of synaesthetic colours might be due to a lack of power, since performance across real and imaginary images is typically lower than for real images (Reddy et al., 2010; Bannert & Bartels, 2013). Indeed, if real differences exist between synaesthetes and controls in the measured BOLD signal, these differences are too small to be detected reliably with sample sizes similar to ours, with no indication about the minimum required sample size. Such a reasoning holds for the average performance, but some subjects did reach performance for colour decoding well above 50%. Yet, the distribution of individual scores were all very similar for controls and synaesthetes (see Figs. 5-6). There was some correlation between the performances of colour (‘Col’) and synaesthetic (‘Syn’) classifiers in retinotopic areas (especially V1), but it was similar in synaesthetes and controls (the differences between synaesthetes and controls for the ‘Syn’ classifier were in fact even weaker when including the ‘Col’ performance as a covariate).

For the statistical analysis, we adopted the “new statistical approach” proposed by (Cumming, 2012) and focussed on confidence intervals of effect sizes instead of the less informative thresholded p-value maps (Hupé, 2015). In order to facilitate the comparison of our study with previous studies, we indicated when the comparisons could be considered as “significant” (a 95% CI not crossing the chance level corresponds to p < 0.05) when correcting the risk level for multiple comparisons. Note however that correction for multiple comparisons corresponds to an ill-posed problem, because there is no unique and objective way to define the family of tests (Hupé, 2015). Such a problem is pretty obvious in our case, where the number of considered ROIs depends on our choice of regrouping or not ROIs, and by how much. We applied a Bonferroni correction over twelve ROIs, but we could have considered the family across the five types of classifiers (so at least 60 tests). However, by focusing on the extent of the CIs, the conclusions do not change much for different levels of CI (the extent of a 99.58% CIs is just a bit larger than for a 95% CI): for all the cases that may suggest differences between groups, the true differences compatible with our observations may be either close to absent (difference close to or including 0, or odds ratio close to or including 1) or at most up to about 15% (or odds ratio = 1.5), a value that one may consider meaningful. As in most studies currently published in cognitive neuroscience dealing with small effects, the width of our confidence intervals is too wide to reach any definitive conclusion on the sole basis of one test (lack of power). Our choice of CI presentation, however, brings useful information allowing cumulative science (Yarkoni, Poldrack, Van Essen, & Wager, 2010) and shows that if any real difference exists, it is probably not very large because corresponding to less than a 15% difference of performance.

## Conclusion

Identifying the neural correlates of the synaesthetic experience of colours may still be beyond the reach of present technology, including hardware (3T MR scanner) and advanced data analysis techniques such as MVPA, and that we still do not find any evidence of common neural coding of real and synaesthetic colours (Hupé et al., 2012b). However, across all our analyses, we did find several “significant” differences for several comparisons, which we listed in the Results section and detailed in Supporting Information, Table S1. Other studies also did report so-called “significant” effects, even though the methods to determine the significance levels were questionable in most studies (Hupé & Dojat, 2015). We applied the latest recommendations for group-level cluster-wise inferences (9-mmm FWHM spatial smoothing, cluster-defining threshold = 0.001, cluster-based pFWE <0.05, groups of 20 participants), yet these criteria do not protect well against false positives (Eklund et al., 2016). In our study, for each “significant” effect, we had a set of independent measures to further explore any difference that may be real: the performance of other classifiers as well as individual differences (the strength of the synaesthetic associations measured in Stoop-like psychophysics tests). We never found any coherence across different measures. Moreover, the locations of the “significant” effects appeared quite randomly across the brain. As long as no other study replicates any of these “differences”, we should keep in mind that they could be false positives. Our study thus further shows that common statistical practices based on Null Hypothesis Significance Tests (NHST) are not adequate for scientific inference (Hupé & Dojat, 2015; Wasserstein & Lazar, 2016; Hupé, 2017). By stressing that we did not find any evidence of common neural coding of real and synaesthetic colours, based on our data as well as past studies, we do not conclude that such a neural coding does not exist. We bring to light what is required to have any chance to reveal the neural bases of the synaesthetic experience using MRI, like more data by subject, higher signal to noise ratio and spatial resolution (e.g., 7 Tesla scanner: Turner, 2016) and maybe larger cohorts. In order to start contributing to this last aim, via the constitution of dedicated data repositories and meta-analyses, our data are freely available on request (https://shanoir.irisa.fr/Shanoir/login.seam, contact M. Dojat). Please refer to the present paper in case of the reuse of these datasets.

## Acknowledgements

Research funded by *Agence Nationale de la Recherche* ANR-11-BSH2-010. Mathieu J. Ruiz was supported by a Ph.D. allowance from the Université Grenoble Alpes. Grenoble MRI facility IRMaGe was partly funded by the French program *Investissement d’avenir* run by the Agence Nationale de la Recherche: grant *Infrastructure d’avenir en Biologie Santé* ANR-11-INBS-0006.

## Appendix

**Supporting Information** is available. It contains:

Table S1. Clusters identified based on whole brain analyses and tested post-hoc with MVPA.

Figure S1. Right occipito-parietal cortex cluster identified based on whole brain univariate analysis

Figure S2. Left anterior insula cluster identified based on whole brain univariate analysis

Figure S3. Right frontal cortex cluster identified based on whole brain univariate analysis

Figure S4. Alternative version of Figure 5, based on mixed-effect generalized linear models

Figure S5. Alternative version of Figure 6, based on mixed-effect generalized linear models

## Author Contributions

M.J.R., M.D, and J.-M.H. designed research; M.J.R. and M.D. performed research; M.J.R., M.D, and J.-M.H. analysed data; J.-M.H. wrote the paper; M.J.R., M.D, and J.-M.H. revised the paper.

## References

Bannert, M. M., & Bartels, A. (2013). Decoding the yellow of a gray banana. Curr. Biol., 23(22), 2268–2272. doi: 10.1016/j.cub.2013.09.016

Bates, D., Mächler, M., Bolker, B., & Walker, S. (2015). Fitting linear mixed-effects models using lme4. J. Stat. Soft., 67(1), 48. doi: 10.18637/jss.v067.i01

Bordier, C., Hupé, J. M., & Dojat, M. (2015). Quantitative evaluation of fMRI retinotopic maps, from V1 to V4, for cognitive experiments. Front. Hum. Neurosci., 9, 277. doi: 10.3389/fnhum.2015.00277

Brouwer, G. J., & Heeger, D. J. (2009). Decoding and reconstructing color from responses in human visual cortex. J. Neurosci., 29(44), 13992–14003. doi: 10.1523/JNEUROSCI.3577-09.2009

Caspers, J., Zilles, K., Eickhoff, S. B., Schleicher, A., Mohlberg, H., & Amunts, K. (2013). Cytoarchitectonical analysis and probabilistic mapping of two extrastriate areas of the human posterior fusiform gyrus. Brain Struct. Funct., 218(2), 511–526. doi: 10.1007/s00429-012-0411-8

Caspers, S., Eickhoff, S. B., Geyer, S., Scheperjans, F., Mohlberg, H., Zilles, K., & Amunts, K. (2008). The human inferior parietal lobule in stereotaxic space. Brain Struct. Funct., 212(6), 481–495. doi: 10.1007/s00429-008-0195-z

Caspers, S., Geyer, S., Schleicher, A., Mohlberg, H., Amunts, K., & Zilles, K. (2006). The human inferior parietal cortex: cytoarchitectonic parcellation and interindividual variability. NeuroImage, 33(2), 430–448. doi: 10.1016/j.neuroimage.2006.06.054

Chiou, R., & Rich, A. N. (2014). The role of conceptual knowledge in understanding synaesthesia: Evaluating contemporary findings from a ‘hub-and-spoke’ perspective. Front. Psychol., 5. doi: 10.3389/fpsyg.2014.00105

Choi, H. J., Zilles, K., Mohlberg, H., Schleicher, A., Fink, G. R., Armstrong, E., & Amunts, K. (2006). Cytoarchitectonic identification and probabilistic mapping of two distinct areas within the anterior ventral bank of the human intraparietal sulcus. J. Comp. Neurol., 495(1), 53–69. doi: 10.1002/cne.20849

Chun, C. A., & Hupé, J. M. (2013). Mirror-touch and ticker tape experiences in synesthesia. Front. Psychol., 4, 776. doi: 10.3389/fpsyg.2013.00776

Chun, C. A., & Hupé, J. M. (2016). Are synesthetes exceptional beyond their synesthetic associations? A systematic comparison of creativity, personality, cognition, and mental imagery in synesthetes and controls. Brit. J. Psychol., 107, 397–418. doi: 10.1111/bjop.12146

Cichy, R. M., Heinzle, J., & Haynes, J. D. (2012). Imagery and perception share cortical representations of content and location. Cereb. Cortex, 22(2), 372–380. doi: 10.1093/cercor/bhr106

Cox, D. D., & Savoy, R. L. (2003). Functional magnetic resonance imaging (fMRI) “brain reading”: detecting and classifying distributed patterns of fMRI activity in human visual cortex. NeuroImage, 19(2 Pt 1), 261–270.

Cumming, G. (2012). Understanding The New Statistics: Effect Sizes, Confidence Intervals, and Meta-Analysis. New York: Routledge.

Dehaene, S., & Cohen, L. (2011). The unique role of the visual word form area in reading. Trends Cogn. Sci., 15(6), 254–262. doi: 10.1016/j.tics.2011.04.003

Dojat, M., Pizzagalli, F., & Hupé, J. M. (2018). Magnetic resonance imaging does not reveal structural alterations in the brain of grapheme-color synesthetes. PLoS ONE, 13(4), e0194422. doi: 10.1371/journal.pone.0194422

Eagleman, D. M., Kagan, A. D., Nelson, S. S., Sagaram, D., & Sarma, A. K. (2007). A standardized test battery for the study of synesthesia. J. Neurosci. Methods, 159(1), 139–145. doi: 10.1016/j.jneumeth.2006.07.012

Edquist, J., Rich, A. N., Brinkman, C., & Mattingley, J. B. (2006). Do synaesthetic colours act as unique features in visual search? Cortex, 42(2), 222–231.

Eickhoff, S. B., Paus, T., Caspers, S., Grosbras, M. H., Evans, A. C., Zilles, K., & Amunts, K. (2007). Assignment of functional activations to probabilistic cytoarchitectonic areas revisited. NeuroImage, 36(3), 511–521. doi: 10.1016/j.neuroimage.2007.03.060

Eklund, A., Nichols, T. E., & Knutsson, H. (2016). Cluster failure: Why fMRI inferences for spatial extent have inflated false-positive rates. Proc. Natl. Acad. Sci. USA, 113(28), 7900–7905. doi: 10.1073/pnas.1602413113

Flournoy, T. (1893). Des Phénomènes de Synopsie (Audition Colorée): Photismes, Schèmes Visuels, Personnifications. Paris: Alcan.

Formisano, E., & Kriegeskorte, N. (2012). Seeing patterns through the hemodynamic veil--the future of pattern-information fMRI. NeuroImage, 62(2), 1249–1256. doi: 10.1016/j.neuroimage.2012.02.078

Galton, F. (1880). Statistics of mental imagery. Mind, 5, 301–318.

Gebuis, T., Nijboer, T. C., & Van der Smagt, M. J. (2009). Multiple dimensions in bi-directional synesthesia. Eur. J. Neurosci., 29(8), 1703–1710. doi: 10.1111/j.1460-9568.2009.06699.x.

Grill-Spector, K., Kourtzi, Z., & Kanwisher, N. (2001). The lateral occipital complex and its role in object recognition. Vision Res., 41(10-11), 1409–1422. doi: 10.1016/S0042-6989(01)00073-6

Hebart, M. N., & Baker, C. I. (2017). Deconstructing multivariate decoding for the study of brain function. NeuroImage. doi: 10.1016/j.neuroimage.2017.08.005

Hupé, J. M. (2015). Statistical inferences under the Null hypothesis: Common mistakes and pitfalls in neuroimaging studies. Front. Neurosci., 9, 18. doi: 10.3389/fnins.2015.00018

Hupé, J. M. (2017). Comment on “Ducklings imprint on the relational concept of ‘same or different’”. Science, 355(6327), 806–806. doi: 10.1126/science.aah6047

Hupé, J. M., Bordier, C., & Dojat, M. (2012a). A BOLD signature of eyeblinks in the visual cortex. NeuroImage, 61, 149–161. doi: 10.1016/j.neuroimage.2012.03.001

Hupé, J. M., Bordier, C., & Dojat, M. (2012b). The neural bases of grapheme-color synesthesia are not localized in real color sensitive areas. Cereb. Cortex, 22, 1622:1633. doi: 10.1093/cercor/bhr236

Hupé, J. M., & Dojat, M. (2015). A critical review of the neuroimaging literature on synesthesia. Front. Hum. Neurosci., 9, 103. doi: 10.3389/fnhum.2015.00103

Janik McErlean, A. B., & Banissy, M. J. (2017). Color processing in synesthesia: what synesthesia can and cannot tell us about mechanisms of color processing. Top. Cogn. Sci., 9(1), 215–227. doi: 10.1111/tops.12237

Kriegeskorte, N., Goebel, R., & Bandettini, P. (2006). Information-based functional brain mapping. Proc. Natl. Acad. Sci. USA, 103(10), 3863–3868. doi: 10.1073/pnas.0600244103

Lorenz, S., Weiner, K. S., Caspers, J., Mohlberg, H., Schleicher, A., Bludau, S., … Amunts, K. (2017). Two new cytoarchitectonic areas on the human mid-fusiform gyrus. Cereb. Cortex, 27(1), 373–385. doi: 10.1093/cercor/bhv225

Mumford, J. A., Turner, B. O., Ashby, F. G., & Poldrack, R. A. (2012). Deconvolving BOLD activation in event-related designs for multivoxel pattern classification analyses. NeuroImage, 59(3), 2636–2643. doi: 10.1016/j.neuroimage.2011.08.076

Norman, K. A., Polyn, S. M., Detre, G. J., & Haxby, J. V. (2006). Beyond mind-reading: multi-voxel pattern analysis of fMRI data. Trends Cogn. Sci., 10(9), 424–430. doi: 10.1016/j.tics.2006.07.005

Parkes, L. M., Marsman, J. B., Oxley, D. C., Goulermas, J. Y., & Wuerger, S. M. (2009). Multivoxel fMRI analysis of color tuning in human primary visual cortex. J. Vis., 9(1), 1 1-13. doi: 10.1167/9.1.1

Pedregosa, F., Varoquaux, G., Gramfort, A., Michel, V., Thirion, B., Grisel, O., … Duchesnay, E. (2011). Scikit-learn: Machine Learning in Python. J. Mach. Learn. Res., 12, 2825–2830.

Poldrack, R. A. (2007). Region of interest analysis for fMRI. Soc. Cogn. Affect. Neurosci., 2(1), 67–70. doi: 10.1093/scan/nsm006

Reddy, L., Tsuchiya, N., & Serre, T. (2010). Reading the mind’s eye: decoding category information during mental imagery. NeuroImage, 50(2), 818–825. doi: 10.1016/j.neuroimage.2009.11.084

Reeder, R. R. (2016). Individual differences shape the content of visual representations. Vision Res. doi: 10.1016/j.visres.2016.08.008

Rouw, R., & Scholte, H. S. (2010). Neural basis of individual differences in synesthetic experiences. J. Neurosci., 30(18), 6205–6213. doi: 10.1523/JNEUROSCI.3444-09.2010

Rouw, R., & Scholte, H. S. (2016). Personality and cognitive profiles of a general synesthetic trait. Neuropsychologia, 88, 35–48. doi: 10.1016/j.neuropsychologia.2016.01.006

Ruiz, M. J., & Hupé, J. M. (2015). Assessment of the hemispheric lateralization of grapheme-color synesthesia with Stroop-type tests. PLoS ONE, 10(3), e0119377. doi: 10.1371/journal.pone.0119377

Ruiz, M. J., Hupé, J. M., & Dojat, M. (2012). Use of pattern-information analysis in vision science: a pragmatic examination. Paper presented at the MLMI, Nice.

Scheperjans, F., Hermann, K., Eickhoff, S. B., Amunts, K., Schleicher, A., & Zilles, K. (2008). Observer-independent cytoarchitectonic mapping of the human superior parietal cortex. Cereb. Cortex, 18(4), 846–867. doi: 10.1093/cercor/bhm116

Simner, J., & Carmichael, D. A. (2015). Is synaesthesia a dominantly female trait? Cogn. Neurosci., 6(2-3), 68–76. doi: 10.1080/17588928.2015.1019441

Simner, J., Mulvenna, C., Sagiv, N., Tsakanikos, E., Witherby, S. A., Fraser, C., … Ward, J. (2006). Synaesthesia: The prevalence of atypical cross-modal experiences. Perception, 35(8), 1024–1033. doi: 10.1068/p5469

Stelzer, J., Lohmann, G., Mueller, K., Buschmann, T., & Turner, R. (2014). Deficient approaches to human neuroimaging. Front. Hum. Neurosci., 8(462). doi: 10.3389/fnhum.2014.00462

Thirion, B., Duchesnay, E., Hubbard, E., Dubois, J., Poline, J. B., Lebihan, D., & Dehaene, S. (2006). Inverse retinotopy: inferring the visual content of images from brain activation patterns. NeuroImage, 33(4), 1104–1116. doi: 10.1016/j.neuroimage.2006.06.062

Tootell, R. B. H., & Nasr, S. (2017). Columnar segregation of magnocellular and parvocellular streams in human extrastriate cortex. J. Neurosci., 37(33), 8014–8032. doi: 10.1523/jneurosci.0690-17.2017

Turner, R. (2016). Uses, misuses, new uses and fundamental limitations of magnetic resonance imaging in cognitive science. Philos. Trans. R. Soc. Lond. B. Biol. Sci., 371(1705). doi: 10.1098/rstb.2015.0349

Vasseur, F., Delon-Martin, C., Bordier, C., Warnking, J., Lamalle, L., Segebarth, C., & Dojat, M. (2010). fMRI retinotopic mapping at 3 T: Benefits gained from correcting the spatial distortions due to static field inhomogeneity. J. Vis., 10 (12), 30. doi: 10.1167/10.12.30

Ward, J. (2013). Synesthesia. Annual Reviews in Psychology, 64, 49–75. doi: 10.1146/annurev-psych-113011-143840

Wasserstein, R. L., & Lazar, N. A. (2016). The ASA’s Statement on p-Values: Context, Process, and Purpose. The American Statistician, 70(2), 129–133. doi: 10.1080/00031305.2016.1154108

Watson, M. R., Chromy, J., Crawford, L., Eagleman, D. M., Enns, J. T., & Akins, K. A. (2017). The prevalence of synaesthesia depends on early language learning. Conscious Cogn., 48, 212–231. doi: 10.1016/j.concog.2016.12.004

Witthoft, N., & Winawer, J. (2013). Learning, memory, and synesthesia. Psychol. Sci., 24(3), 258–265. doi: 10.1177/0956797612452573

Yarkoni, T., Poldrack, R. A., Van Essen, D. C., & Wager, T. D. (2010). Cognitive neuroscience 2.0: building a cumulative science of human brain function. Trends Cogn. Sci., 14(11), 489–496. doi: 10.1016/j.tics.2010.08.004

