## Supporting Information for "Multivariate pattern analysis of fMRI data for imaginary and real colours in grapheme-colour synaesthesia"

**Table S1. Clusters identified based on whole brain analyses and tested *post-hoc* with MVPA.**

**Figure S1. Right occipito-parietal cortex cluster identified based on whole brain univariate analysis**

**Figure S2. Left anterior insula cluster identified based on whole brain univariate analysis**

**Figure S3. Right frontal cortex cluster identified based on whole brain univariate analysis**

**Figure S4. Alternative version of Figure 5, based on mixed-effect generalized linear models**

**Figure S5. Alternative version of Figure 6, based on mixed-effect generalized linear models**

**Table S1. Clusters identified based on whole brain analyses and tested *post-hoc* with MVPA.**

| Whole brain method to identify clusters | | | | | | | | MVPA tests in post-hoc clusters | | | | | | | | | | |
|---|---|---|---|---|---|---|---|---|---|---|---|---|---|---|---|---|---|---|
| | | | | | | | | | Col | | Syn | | C2S | | S2C | | g1g2 | |
| analysis | stimuli | stat | comparison | contrast | size | MNI XYZ | name | | S>C | S>0.25 | S>C | S>0.25 | S>C | S>0.25 | S>C | S>0.25 | S>C | S>0.25 |
| **univariate** | graph. | T | P-s | Syn>Con | 314 | [-60 -4 21] | left precentral | yes | | | | | | | | | | |
| | | T | P-s & 2-s | Con>Syn | | | | | | | | | | | | | | |
| | | F | P-s & 2-s | Syn>Con | | | | | | | | | | | | | | |
| ***Fig. S1*** | | F | P-s & 2-s | Con>Syn | 388 | [33 -70 33] | right occipital-parietal | yes | 0.011 | 0.003 | 0.016 | 0.122 | | | | | | |
| | | | P-s | Con>Syn | 361 | [-24 21 -8] | left insula | yes | | | | | | | | | | |
| | colours | T | P-s | Syn>Con | 327 | [-39 -16 -8] | left posterior insula | yes | | | | | | | | | | |
| ***Fig. S2*** | | | P-s | Syn>Con | 203 | [-35 35 -3] | left anterior insula | yes | | | | | | | 0.029 | 0.077 | | |
| | | | P-s | Syn>Con | 142 | [-32 -16 -33] | left parahippocampal | yes | | | | | | | | | | |
| | | | 2-s | Syn>Con | 513 | [47 -61 6] | right middle temporal | yes | | | | | | | | | | |
| ***Fig. S3*** | | | 2-s | Syn>Con | 385 | [5 30 42] | right superior, frontal | yes | | | 0.023 | 0.087 | | | | | | |
| | | T | P-s & 2-s | Con>Syn | | | | | | | | | | | | | | |
| | | F | P-s & 2-s | Syn>Con | | | | | | | | | | | | | | |
| | | F | P-s | Con>Syn | 128 | [-17 18 21] | white matter | no | | | | | | | | | | |
| **MVPA** | colours | Col | P-s & 2-s | Syn>Con | | | | | | | | | | | | | | |
| | | Col | P-s & 2-s | Con>Syn | | | | | | | | | | | | | | |
| | graph. | Syn | P-s | Syn>Con | 459 | [24 -43 44] | right parietal | yes | | | $4.10^{-7}$ | | | | | | | |
| | | | 2-s | Syn>Con | 351 | [-24 -40 53] | left parietal | yes | | | $2.10^{-4}$ | | | | | | | |
| | | Syn | 1-s | Syn>0.25 | | | | | | | | | | | | | | |
| | | Syn | P-s & 2-s | Con>Syn | | | | | | | | | | | | | | |
| | | Syn | 1-s | Con>0.25 | | | | | | | | | | | | | | |
| | all | C2S | P-s & 2-s | Syn>Con | | | | | | | | | | | | | | |
| | | C2S | 1-s | Syn>0.25 | | | | | | | | | | | | | | |
| | | C2S | P-s & 2-s | Con>Syn | | | | | | | | | | | | | | |
| | | C2S | 1-s | Con>0.25 | | | | | | | | | | | | | | |
| | all | S2C | P-s | Syn>Con | 297 | [39 -70 2] | right occipito-temporal | yes | | | | | | | $7.10^{-7}$ | $1.10^{-5}$ | | |
| | | | 2-s | Syn>Con | 459 | [39 -73 5] | right occipito-temporal | yes | | | | | | | $6.10^{-6}$ | $7.10^{-6}$ | | |
| | | | 2-s | Syn>Con | 648 | [-27 -1 -7] | left putamen | yes | | | | | | | $3.10^{-4}$ | $7.10^{-4}$ | | |
| | | S2C | 1-s | Syn>0.25 | 486 | [42 -73 2] | right occipito-temporal | yes | | | | | | | $9.10^{-6}$ | $5.10^{-6}$ | | |
| ***Fig. 8*** | | | 1-s | Syn>0.25 | 729 | [-33 -28 50] | left parietal | yes | | | 0.012 | 0.004 | | | $5.10^{-4}$ | $2.10^{-6}$ | | |
| | | S2C | P-s & 2-s | Con>Syn | | | | | | | | | | | | | | |
| | | S2C | 1-s | Con>0.25 | | | | | | | | | | | | | | |
| | graph. | g1g2 | P-s & 2-s | Syn>Con | | | | | | | | | | | | | | |
| | | g1g2 | 1-s | Syn>0.25 | | | | | | | | | | | | | | |
| | | g1g2 | P-s & 2-s | Con>Syn | 837 | [-42 20 26] | left inferior frontal | no | | | | | | | | | | |
| | | g1g2 | 1-s | Con>0.25 | | | | | | | | | | | | | | |

Clusters potentially involved in synaesthesia were identified based on whole brain univariate analysis and searchlight MVPA. For each *analysis* the line in the table indicates which *stimuli* were presented ('graph.': achromatic letters and digits; 'all': MVPA based on both graphemes and coloured rings), which statistics (*stat*) was used to create individual whole brain maps (first-level analysis), the statistical test (*comparison:* P-s = paired-sample *T*-test; 2-s = two-sample *T*-test; 1-s = one-sample *T*-test) performed for the second level analysis as well as the statistical *contrast*. For all individual statistical maps (first level analysis), we applied a spatial smoothing with FWHM = 9 mm for univariate analyses and no smoothing for MVPA. For second-level analyses, the cluster forming threshold was set at $p = 0.001$. We list all clusters significant at $p$FWE < 0.05, their *size* in $mm^3$ (voxel size was 1.5 $mm^3$ for univariate analyses and 3 $mm^3$ for multivariate analyses), the coordinates in the *MNI* space of the voxel with the smallest *p*-value in the cluster as well as the *name* used in the main text, corresponding to their approximate location. Empty lines mean the absence of any significant cluster. Grey font was used for statistical contrasts for which we did not have any reason to expect any difference. The right part of the table lists the comparisons of MVPA scores within these *post-hoc* clusters. The names of the MVPA classifiers are explained in Figure 3. We compared the scores of synaesthetes and controls with paired *T*-tests and report the *p*-values that were below 0.05 (two-sided tests, not corrected for multiple comparisons; the results of two-sample *T*-tests were similar). The scores of controls were never significantly larger than the scores of synaesthetes. We also tested the scores of synaesthetes against chance (two-sided one-sample tests) and reported *p*-values systematically when there was a difference between synaesthetes and controls. For the results of MVPA tests in clusters defined by the whole brain MVPA searchlight, we shaded in grey the cells corresponding to circular analysis. Note that for the results of the 'S2C' classifier in clusters based on 'S2C', the comparison of synaesthetes and controls and the comparison of synaesthetes against chance are not independent (the scores of controls in these cluster were in fact on average below chance).

**Figure S1. Right occipito-parietal cortex cluster identified based on whole brain univariate analysis**

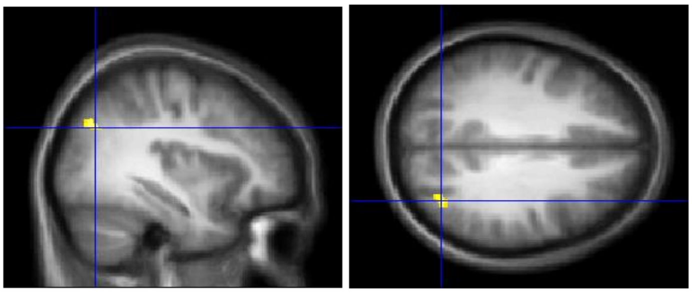

The univariate analysis of *F*-contrast for achromatic graphemes revealed a significant cluster (*p*FWE < 0.05) in the right occipito-parietal cortex (MNI XYZ = [33 -70 33], k = 111) for the contrast Con>Syn (paired *T*-test). MVPA tests in this cluster revealed that synaesthetes decoded graphemes better based on training on graphemes (‘Syn’ classifier, 95% CI of the difference = [1.4 11.3]%). The performance of synaesthetes was also slightly above chance (95% CI = [24 31]%) but did not correlate with photism strength (p = 0.67). (This result is paradoxical since the modulation by graphemes was higher in Controls – that’s how the ROI was defined – so differences of BOLD signals could have favoured the ‘Syn’ classifier for controls). In this cluster, synaesthetes also decoded colours better based on training on colours (‘Col’ classifier, 95% CI of the difference = [0.9 6.1]%). The performance of synaesthetes was also significantly above chance (95% CI = [26 30]%) but did not correlate with photism strength (*p* = 0.66).

**Figure S2. Left anterior insula cluster identified based on whole brain univariate analysis**

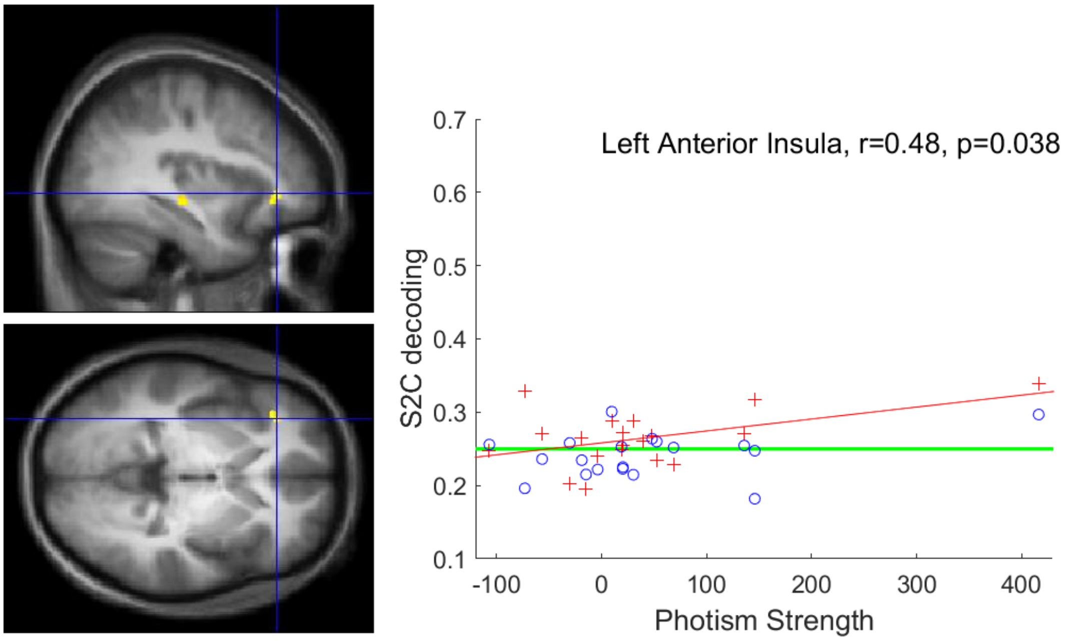


The univariate analysis of *T*-contrast for colour rings revealed a significant cluster (*p*FWE < 0.05) in the left anterior insula (MNI XYZ = [-35 35 -3], k = 60) for the contrast Syn>Con (paired *T*-test). MVPA tests in this cluster revealed that synaesthetes decoded colours better based on training on graphemes (‘S2C’ classifier, 95% CI of the difference = [0.3 4.6]%). The performance of synaesthetes was also slightly above chance (95% CI = [25 28]%) and slightly correlated with the strength of synaesthetic associations (same conventions as in Figure 7). However, the correlation is driven by only one data point (non-parametric Spearman test on ranks, *p* = 0.30).

**Figure S3. Right frontal cortex cluster identified based on whole brain univariate analysis**

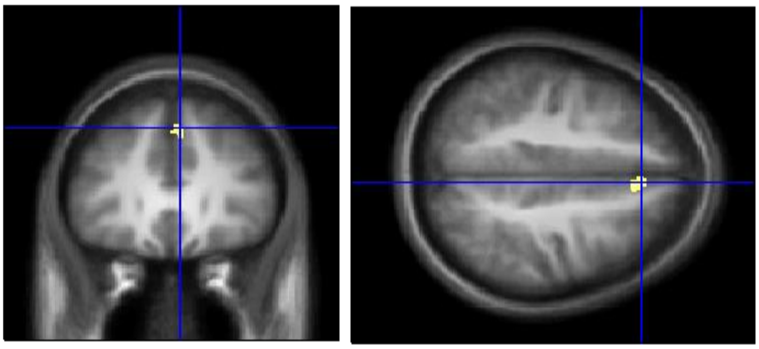

The univariate analysis of *T*-contrast for colour rings revealed a significant cluster (*p*FWE < 0.05) in the right frontal cortex (MNI XYZ = [5 30 42], k = 114) for the contrast Syn>Con (two-sample *T*-test). MVPA tests in this cluster revealed that synaesthetes decoded graphemes better based on training on graphemes ('Syn' classifier, 95% CI of the difference = [1 12]%). The performance of synaesthetes was also slightly above chance (95% CI = [24 34]%) but did not correlate with photism strength (*p* = 0.54).

**Figure S4. Alternative version of Figure 5, based on mixed-effect generalized linear models**

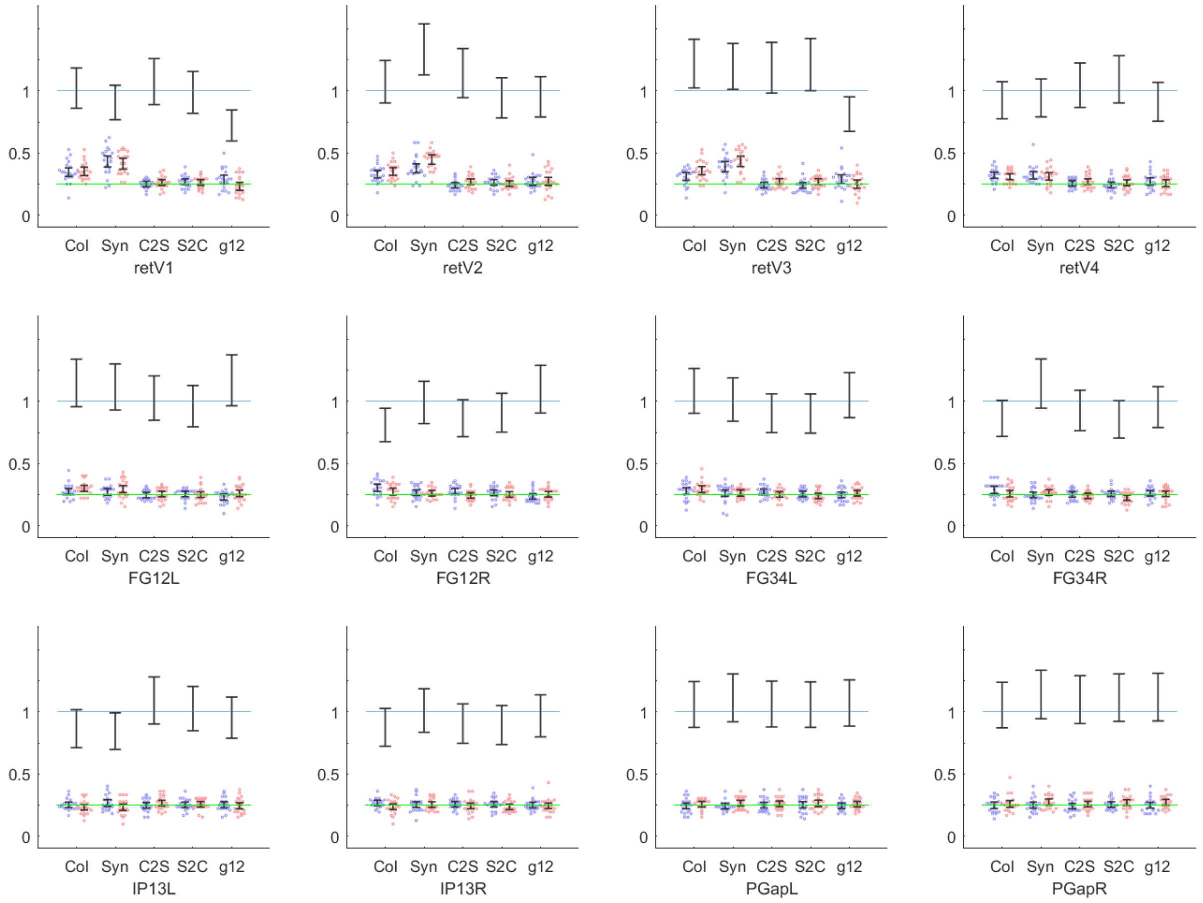


Here the difference of performance between synaesthetes and controls was estimated by a mixed-effect generalized linear models with a binomial family and a logit link function. The y-axis represents therefore not only the performance of classifiers for individual subjects and their group average and CI like in Figure 5, but also the odd-ratio of synaesthetes against their matched controls (1 = no difference between groups, blue line; whiskers denote 95% CI). Estimation is slightly more precise with this more powerful analysis.

**Figure S5. Alternative version of Figure 6, based on mixed-effect generalized linear models**

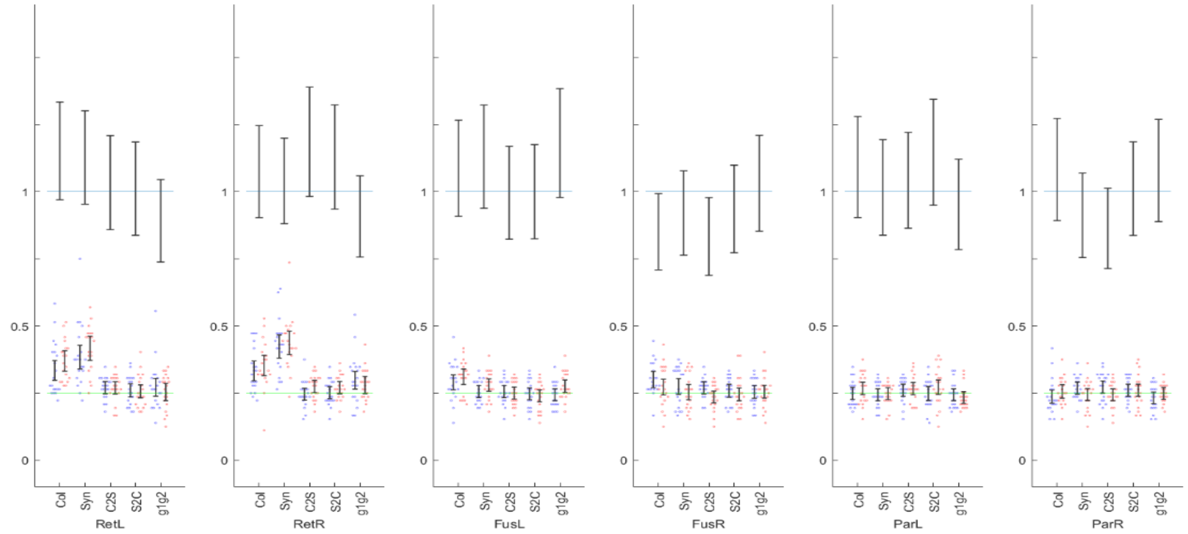
